# Design of facilitated dissociation enables control over cytokine signaling duration

**DOI:** 10.1101/2024.11.15.623900

**Authors:** Adam J. Broerman, Christoph Pollmann, Mauriz A. Lichtenstein, Mark D. Jackson, Maxx H. Tessmer, Won Hee Ryu, Mohamad H. Abedi, Danny D. Sahtoe, Aza Allen, Alex Kang, Joshmyn De La Cruz, Evans Brackenbrough, Banumathi Sankaran, Asim K. Bera, Daniel M. Zuckerman, Stefan Stoll, Florian Praetorius, Jacob Piehler, David Baker

## Abstract

Protein design has focused primarily on the design of ground states, ensuring they are sufficiently low energy to be highly populated^1^. Designing the kinetics and dynamics of a system requires, in addition, the design of excited states that are traversed in transitions from one low-lying state to another^2,3^. This is a challenging task as such states must be sufficiently strained to be poorly populated, but not so strained that they are not populated at all, and because protein design methods have generally focused on creating near-ideal structures^4–7^. Here we describe a general approach for designing systems which use an induced-fit power stroke^8^ to generate a structurally frustrated^9^ and strained excited state, allosterically driving protein complex dissociation. X-ray crystallography, double electron-electron resonance spectroscopy, and kinetic binding measurements demonstrate that incorporating excited states enables design of effector-induced increases in dissociation rates as high as 6000-fold. We highlight the power of this approach by designing cytokine mimics which can be dissociated within seconds from their receptors.

## Main

Protein-protein interactions orchestrate much of biological function. High affinity interactions enable protein circuits to respond to low concentrations of stimuli and to potently act on targets; fast exchange enables them to respond quickly to changes in stimuli. These two properties usually cannot simultaneously be achieved in binary interactions because they depend on the interaction off-rate in opposite ways: high affinity usually requires a slow dissociation (low off-rate) whereas rapid exchange requires a fast dissociation (high off-rate) (Fig. S1). Several natural systems exhibit “facilitated dissociation”^10–14^ in which a host (A) can bind a partner (B), and an effector (C) can bind to the host-partner complex (AB) to form an excited ternary complex (ABC) from which the partner dissociates quickly (Fig. 1a). In such a system, the partner can bind tightly to the host yet can also be rapidly released by adding the effector^15^. In engineered DNA systems, the kinetic control afforded by an analogous phenomenon (toehold-mediated strand displacement) has enabled the construction of many complex functions^16,17^, but DNA systems have limited utility for directly interfacing with biology. Protein binding/unbinding can be readily coupled to biological processes, but there has been no general approach to design kinetic control over protein interactions.

**Fig. 1.**
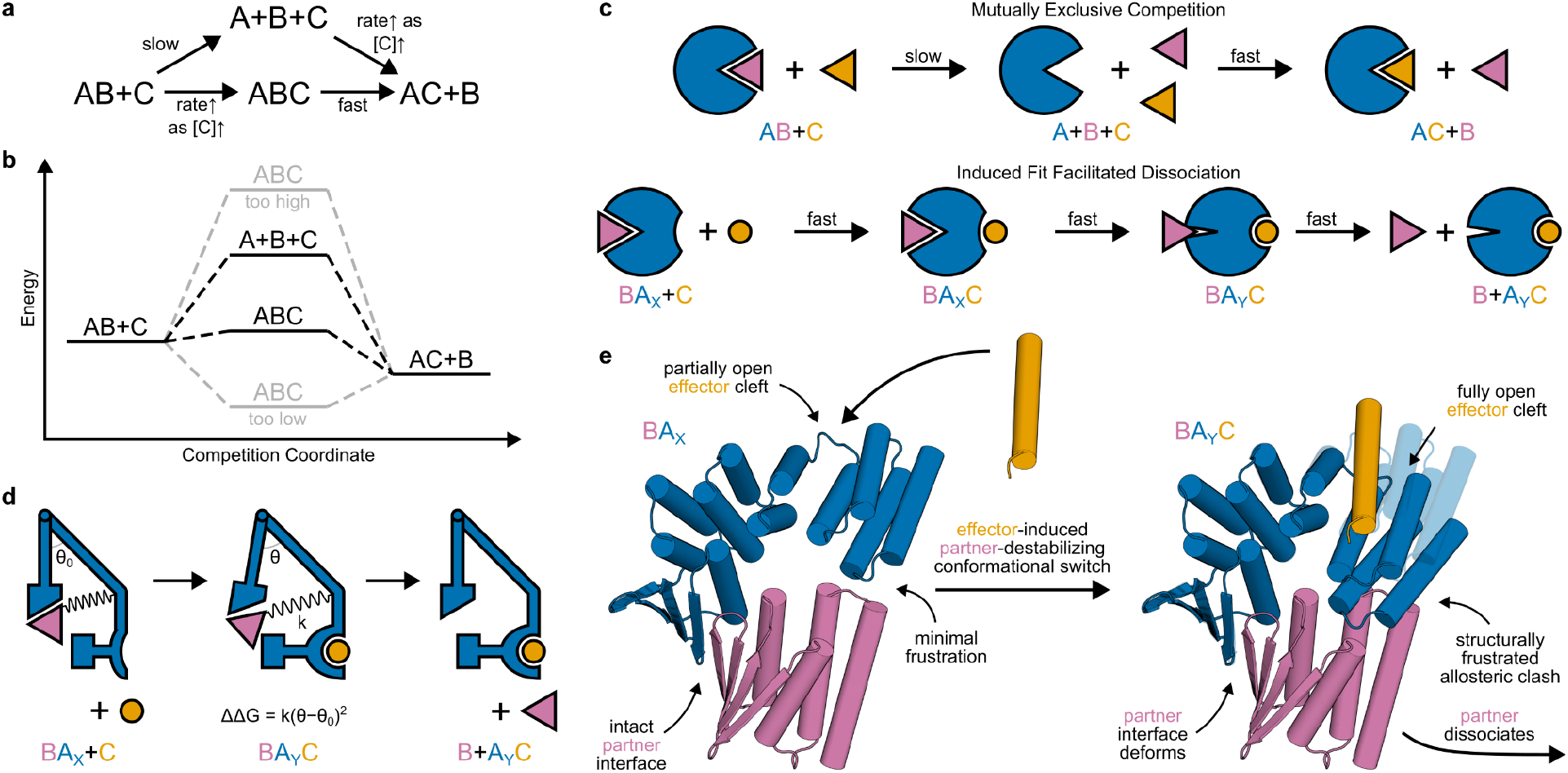
Strategy for designing proteins which reconfigure through facilitated dissociation. **a, b, c**, A high affinity interaction can rapidly exchange through facilitated dissociation (lower pathways), but not through mutually exclusive competition (upper pathways). **a**, Reaction diagram. **b**, Energy diagram. **c**, Schematic of induced-fit facilitated dissociation (bottom) compared to slow mutually exclusive competition (top). The host protein (A, subscripted by conformational state X or Y) is shown in blue, the partner (B) in pink, and the effector (C) in orange. **d**, Schematic of structural frustration resolved through strain in a facilitated dissociation pathway. Hooke’s law can relate the energy of the ternary intermediate to mechanical properties of the protein. **e**, Examples of design models of proteins designed to undergo a facilitated dissociation process starting from a tightly interacting state X (left) through a structurally frustrated ternary intermediate in state Y (right, solid) aligned to state X (transparent) to show the conformational change.

We set out to design protein systems which undergo facilitated dissociation. To bypass the slow step of the partner dissociating spontaneously, the effector must bind to the host-partner complex to form a ternary complex intermediate^18,19^ in which the structure^11,20^ or dynamics^21–23^ of the host-partner interaction are altered to favor dissociation (Fig. 1a); binding the effector must reshape the energy landscape to provide a faster pathway for partner dissociation (Fig. 1b). Given an interaction between a protein binder and partner, we reasoned that host proteins supporting facilitated dissociation could be constructed by rigidly fusing an effector-responsive conformational switch to the binder such that, due to a change in steric clashes, the interaction energy depends on the state of the switch. This would allosterically couple the partner and the effector, and rapid partner dissociation could proceed through a mechanically strained and excited ternary complex intermediate (Fig. 1c). Critically, the energy of this ternary intermediate must be neither too high (otherwise the facilitated dissociation pathway would not be faster) nor too low (otherwise the partner would not dissociate) (Fig. 1b). Treating the ternary complex as a spring using Hooke’s law (Fig. 1d), we reasoned that by varying the geometry and structure of the fusion, we could control the strain and stiffness (spring constant) of the ternary complex to set its energy within this optimal range (Fig. S2).

To effectively access such a strained ternary complex to modulate partner binding, we reasoned the switch should satisfy several criteria. First, the conformational change should be large to enable steric modulation of partner binding affinity. Second, the switch should bind the effector tightly so there is sufficient driving force to expel the partner. Third, the switch should be stiff so that in the strained ternary complex, the strain localizes to the partner interface. Fourth, akin to a power stroke^8^, the effector should rapidly drive the conformational change even against the resisting force associated with forming the strained ternary complex. Design of stimulus-responsive conformational change has been achieved with hinge-like proteins which switch from a closed state (X) to an open state (Y) to bind an effector through a conformational selection mechanism^24^. The rigid-body motion of these hinges can be large (criterion #1) yet rapid because it requires breaking few interactions^24^, the effectors bind tightly (criterion #2) by forming a large, shape complementary interaction in the state Y cleft^24^, and their topology (in which each helix contacts several others, leading to a high packing density) could be stiff^25^ (criterion #3). But since the hinge conformational change must occur before effector binding, it cannot be a driven power stroke^8,26^ (criterion #4). An induced-fit mechanism would be better suited for rapidly driving a large conformational change against a resisting force because effector binding would not be rate-limited by a conformational pre-equilibrium slowed by the resisting force^27,28^.

Our approach to constructing a facilitated dissociation system thus requires designing an induced-fit conformational switch satisfying all these criteria and a switch-binder fusion with working allosteric coupling. Uncoupling the combinatorial problem of simultaneously obtaining both functions, we first used a previously designed hinge (cs221^24^) to test the concept of allosteric coupling via switching steric clashes. For a model partner-binder interaction, we chose a designed heterodimer pair (LHD101A-LHD101B^29^, modified as described in Fig. S3). To couple the effector-induced conformational switch to dissociation of a partner, we generated structured fusions of cs221 and LHD101B such that in state X, the partner LHD101An1 can bind, but in the effector-bound state Y, it would strongly clash (Fig. S4a). We obtained synthetic genes encoding 12 designs, expressed and purified the proteins from *E. coli*, and found that several showed slow and significantly reduced effector association in the presence of the partner (Fig. S4d), indicating the desired allosteric coupling but also the need for designs with a faster, driven pathway for effector binding against the partner (criterion #4).

To design induced-fit effector binding, we reasoned we could design a rigid-body conformational switch in which the effector can make interactions with both conformational states, weakly in one and tightly in the other, and can remain bound throughout the switching transition. Starting from the best cs221-LHD101B fusion design, “allosteric switch 0” AS0, we designed switches with the same effector-bound state Y backbone but with a new state X which retains an open effector binding cleft where the effector could weakly associate (Fig. 1e). We constructed this new state X by shifting the two domains from state Y relative to each other by one helix heptad, then refining the dock between domains (Fig. S5a, methods). This approach maintains the open cleft by introducing minimal rotation and simplifies the multi-state sequence design challenge by minimizing the local structural differences between the two states. Switching between conformational states would occur by a register shift, throughout which the effector could remain bound (Fig. S5b). An effector that is flexible or folds upon binding (and hence does not clash when binding is initiated with state X) could reduce the energy barrier for this transition, enabling the conformational change to more rapidly and efficiently exert force to accelerate partner dissociation^26,30^. Since the new sliding switches retain the state Y backbone validated to clash in AS0, we should observe allosteric coupling if the switch works as designed (Fig. 1e).

We obtained synthetic genes encoding 10 such designs, and found that 4 tightly bound the effector (Fig. S6). To assess facilitated dissociation, we used surface plasmon resonance (SPR) to measure partner-host dissociation kinetics upon exposure to the effector. For these four designs, the partner dissociates slowly in the absence of the effector; addition of effector dramatically increases the rate of dissociation in a concentration-dependent manner (Fig. 2a, S7), but minimally affects the partner off-rate from a control static binder fusion lacking an effector-binding cleft (Fig. S7, LHD101B4^29^). As the effector concentration increases, the effective rate of partner dissociation approaches the independently measured off-rate of the partner from the ternary complex (Fig. 2g, S7), strongly suggesting that this state is an intermediate in the overall facilitated dissociation process (Fig. 2b, lower pathways).

**Fig. 2.**
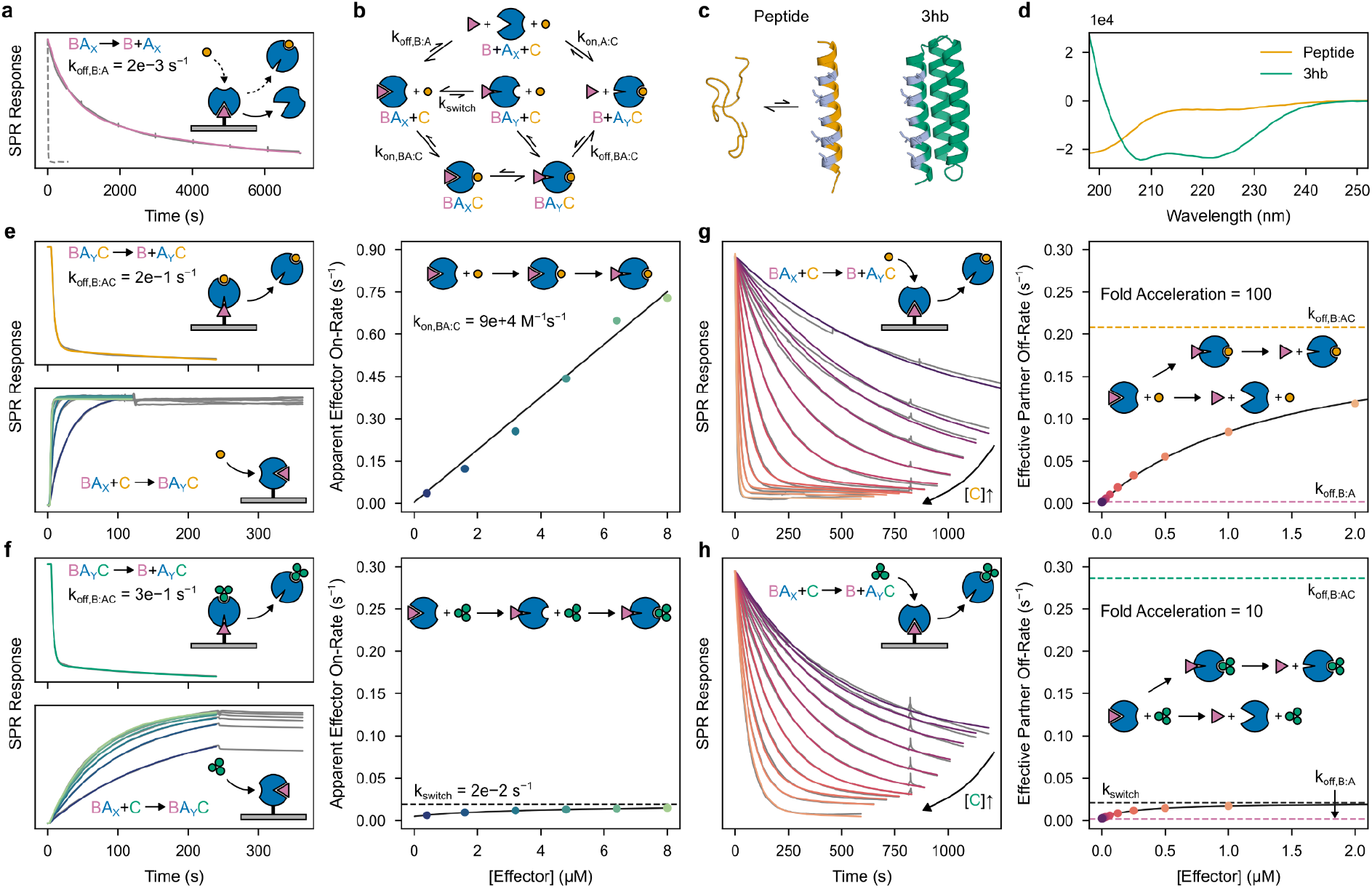
Kinetic characterization of facilitated dissociation in AS1. **a**, Slow dissociation of the partner from the host in the absence of effector (solid) and fast dissociation in the presence of effector (dashed) as assessed by SPR. Slow dissociation data (solid gray) fit with a double exponential (pink). **b**, Kinetic model describing pathways of competition: (top) mutually exclusive competition, (middle) facilitated dissociation with effector binding rate-limited by conformational selection, (bottom) facilitated dissociation with induced-fit effector binding. k_off,B:A_, k_on,BA:C_, k_switch_, and k_off,BA:C_ are rate constants. **c**, Cartoon representations of the peptide (left) and 3hb (right) effectors; interface residues shown in gray. **d**, Circular dichroism (CD) spectra of the peptide and 3hb effectors. **e** and **f**, Kinetic characterization of the formation and breakage of the ternary complex intermediate with the peptide (e) and 3hb (f) effectors. (Top left) fast dissociation of the partner from the ternary complex, data (gray) fit with double exponentials (orange/green); (bottom left) effector association to form the ternary complex and extremely slow subsequent dissociation, data (gray) fit with single exponentials (colors) in the association phase; (right) apparent effector on-rates plotted against effector concentration (circles) and a linear (e) or hyperbolic (f) fit. **g** and **h**, Kinetic characterization of the full facilitated dissociation pathway with the peptide (g) and 3hb (h) effectors. (Left) effector concentration–dependent dissociation of the partner upon addition of effector, data (gray) fit (colors) as described in methods; (right) effective partner off-rates plotted against effector concentration (circles) and fit with a hyperbolic equation (black line). There is a discrepancy in the EC_50_ between this partner dissociation experiment and the 3hb association experiment; this could result from the fusion tag used to affix AS1 to the surface competing with the 3hb for binding the cleft, reducing the apparent 3hb on-rate. In **a** and the left plots of **e**–**h**, cartoons show the arrangement of proteins relative to the SPR chip (gray). In the right plots of **e**–**h**, cartoons show the mechanism that can be inferred from the data.

To further probe the mechanism of facilitated dissociation, we focused on AS1, the design with the tightest effector binding, and characterized two different effectors: the peptide and a three-helix bundle “3hb” (3hb21^24^) (Fig. 2c). The two effectors make nearly identical interactions with AS1, but when unbound, the peptide is disordered and the 3hb is structured (Fig. 2d). The apparent on-rate for binding to the partner-AS1 complex increases linearly with concentration for the peptide (Fig. 2e) but hyperbolically for the 3hb, saturating at the rate of a concentration-independent step (Fig. 2f). Notably, with the 3hb, the effective rate of partner dissociation saturates at this same value (Fig. 2h). The simplest explanation of these results is that the rigid 3hb can only bind to the fully open state Y of AS1 and that the BA_X_→BA_Y_ conformational change is slow (due to partial blocking by the bound partner), so it becomes rate limiting for both the association with the partner-AS1 complex and the overall facilitated dissociation process (Fig. 2b, middle pathway). In contrast to the 3hb, peptide effector binding (Fig. 2e) and resulting partner destabilization (Fig. 2g) both can occur more rapidly than the BA_X_→BA_Y_ conformational change, suggesting that the more flexible peptide effector can bind to AS1 in state X to accelerate this conformational change through an induced-fit mechanism (Fig. 2b, bottom pathway)^30,31^.

We used x-ray crystallography to structurally characterize the different states of our designed systems. For both the AS1 and AS5 systems, the crystal structures of the hosts alone (Fig. 3a, Extended Data Fig. 1a) and of the host-effector complexes (Fig. 3b, Extended Data Fig. 1b) closely match the design models (maximum 1.3 Å Cα RMSD). The unbound structures show the new state X with its open hydrophobic cleft poised for binding, and the effector-bound structures show that binding the effector causes the two switch domains to register shift into the designed state Y and that the binder fusion positions the partner to clash in this state. The structures also show sequence features present in almost all designs (insets in Fig. 3a,b and Extended Data Fig. 1a,b); a positively charged residue (K60) holds the cleft open in state X and moves to interact with the negative dipole of the effector helix in state Y, and the interface between domains in the designed one-heptad register shift contains sequence motifs repeated a heptad apart (on the C-terminal domain, L84 in state X and L91 in state Y pack against the same location on the N-terminal domain).

**Fig. 3.**
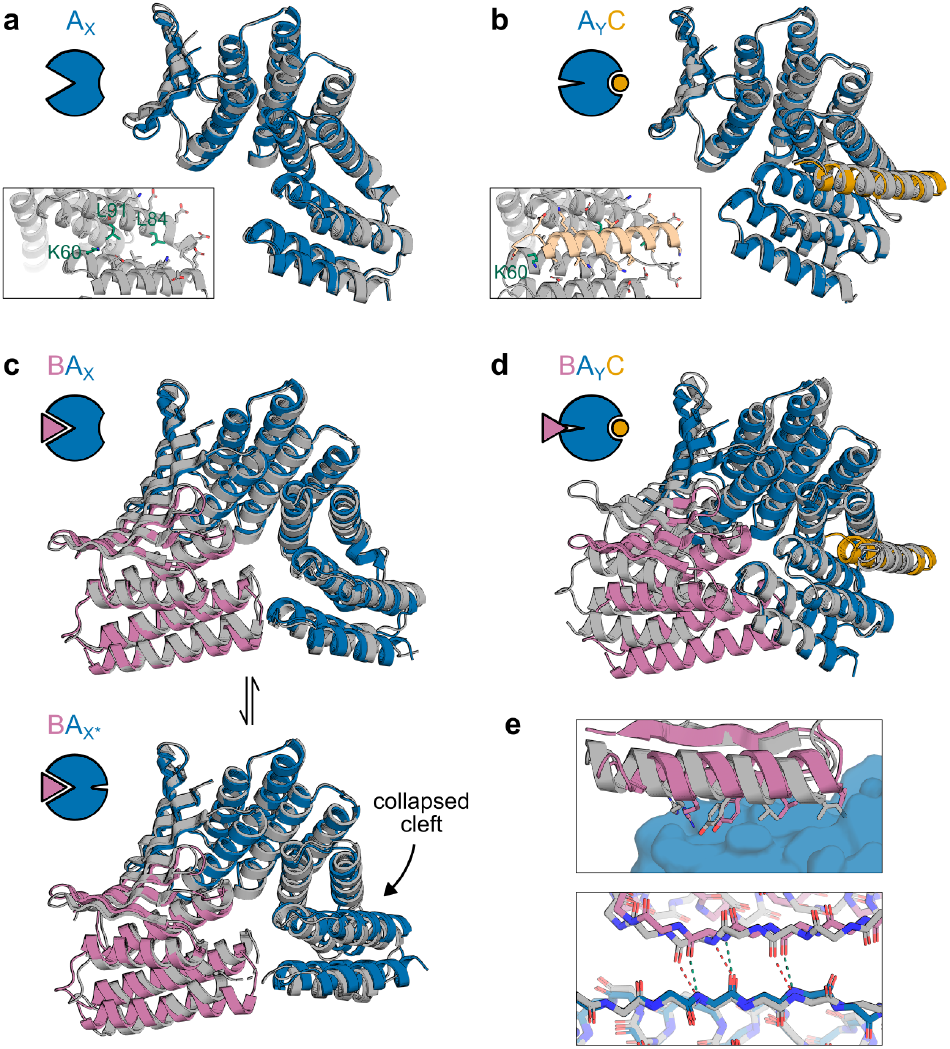
Structural characterization of AS1. **a**, Crystal structure of AS1 alone (gray) overlaid with the design model of AS1 in state X (blue). Inset shows a detailed view of side chains in the partially-open effector-binding cleft. **b**, Cocrystal structure of AS1 and peptide effector (gray) overlaid with the design model of the AS1-effector complex in state Y (AS1 in blue, effector in orange). Inset shows the same view of the side chains in the effector-binding cleft as in (a). **c**, (Top) cocrystal structure of AS1 (with intact cleft) and partner with methylated lysines (gray) overlaid with the design model of the partner-AS1 complex in state X (AS1 in blue, partner in pink). (Bottom) cocrystal structure of AS1 (with collapsed cleft) and partner with methylated lysines (gray) overlaid with the design model of the partner-AS0 complex which adopts a state resembling state X* (AS0 in blue, partner in pink). **d**, Cocrystal structure of the AS1, partner, and peptide effector with methylated lysines (gray) overlaid at the switch region with the design model of the AS1-effector complex in state Y (AS1 in blue, effector in orange) and design model of the partner (pink) aligned to its binding site on the AS1 design model to show the clash. **e**, (Top) detailed view of the partner interface side chains in the ternary complex (gray) and the partner-AS1 complex (pink) interacting with AS1 (blue). (Bottom) detailed view of the backbone hydrogen bonding in the interfacial strand pairing. The partner-AS1 complex (pink and blue) contains unstrained hydrogen bonds (green); the ternary complex (gray) contains strained hydrogen bonds (red).

We solved structures of AS1 bound to the partner from two different crystals, each containing two copies of the complex. Both copies in one crystal and one copy in the other closely match the state X design model with slight fluctuation in the partner binding conformation (Fig. 3c, top; Extended Data Fig. 1d), whereas in the remaining copy, AS1 adopts a different conformation, state X*, in which the effector-binding cleft has collapsed to resemble the closed state X of the original hinge (Fig. 3c, bottom). The new state X of AS1 still slightly clashes with the partner (Extended Data Fig. 1c), likely causing this dynamic partner binding and cleft collapse.

We next sought to determine how the strain in the partner-host-effector ternary complex resolves structurally. At the concentrations of the components used in the above experiments, the AS1 ternary complex is an only transiently populated excited state, but at high concentrations it becomes the dominant state (Fig. S9). This enabled us to solve a crystal structure of AS1 in the ternary complex intermediate with both partner and effector. The switch region closely matches the state Y design model while the rest of the structure strains to resolve the structural frustration from simultaneously binding the partner and occupying state Y (Fig. 3d). This strain localizes to multiple locations in the structure: the binder fusion bends, the portion of the partner which directly clashes with the switch is not resolved in the structure and likely is disordered, and the partner twists at its interface, disrupting the interfacial hydrophobic packing (Fig. 3e, top) and shearing the interfacial strand pairing (Fig. 3e, bottom). Overall, the partner and binder deform from their binary complex conformation much more than the switch (4 Å vs 1 Å Cα RMSD), indicating that the switch is stiff and thus can effectively localize strain to the partner interface to increase its off-rate.

We further characterized the dynamics within the ternary complex using molecular dynamics (MD) simulations and double electron-electron resonance (DEER) spectroscopy to measure pairwise distance distributions between residues on opposing sides of the conformational switch. For the three low-energy states, the AS1 alone, partner-AS1 complex, and AS1-effector complex DEER distance distributions are consistent with the distances between spin labels simulated based on the corresponding crystal structures (Extended Data Fig. 2). For the higher-energy ternary complex, the DEER distance distribution contained two peaks: one aligned with the distances simulated from the ternary complex crystal structure and from two MD trajectories started from the AlphaFold2 (AF2) prediction of the ternary complex, and the other aligned with the distance simulated from a third trajectory (Extended Data Fig. 3). Thus, to resolve strain, the ternary complex can likely adopt multiple conformations with varying amounts of switch deformation, partner deformation, and unfolding of the clashing partner loop (Extended Data Fig. 3).

### Modulation of dissociation rate enhancement

We next explored the factors contributing to the magnitude of the dissociation rate enhancement and the extent to which this could be tuned. Some of the minor strain in the partner-AS1 complex may localize to the partner interface, and indeed the partner dissociates 20-fold faster from AS1 than from an unhindered binder fusion (Fig. S7). Considering the ternary complex as a spring, we hypothesized that increasing the magnitude of strain or deforming in a direction of higher stiffness could increase the energy of the ternary complex intermediate, further enhancing the partner off-rate. To explore this concept, we sought to reposition the partner relative to the conformational switch to minimize strain in the partner-host complex and to maximize strain in the ternary complex while pushing the partner in a more destabilizing direction. Avoiding any clash in state X while maintaining a strong clash in state Y, we sampled a variety of partner positions relative to the switch from AS1, then rebuilt fusions between the switch and the corresponding new positions of the partner binder. AF2 recapitulates the strained AS1 ternary complex with high accuracy (1.0 Å Cα RMSD from the partner-AS1-effector crystal structure), so we used it to select a set of these AS1 variants with substantial deformations spanning a variety of directions. These changes to the fusion region, though distant from both binding sites, caused dramatic variation in the kinetics of partner dissociation and its modulation by the effector (Fig. S10). Most of these variants showed reduced partner off-rates (compared to AS1) in the absence of effector which increased in the presence of effector (Fig. S10), and for several “fast variants” (AS116, AS117, and AS118), adding effector accelerated partner dissociation by up to 2000-fold, reaching rates comparable to that of original AS1 (Fig. 4a, S10).

**Fig. 4.**
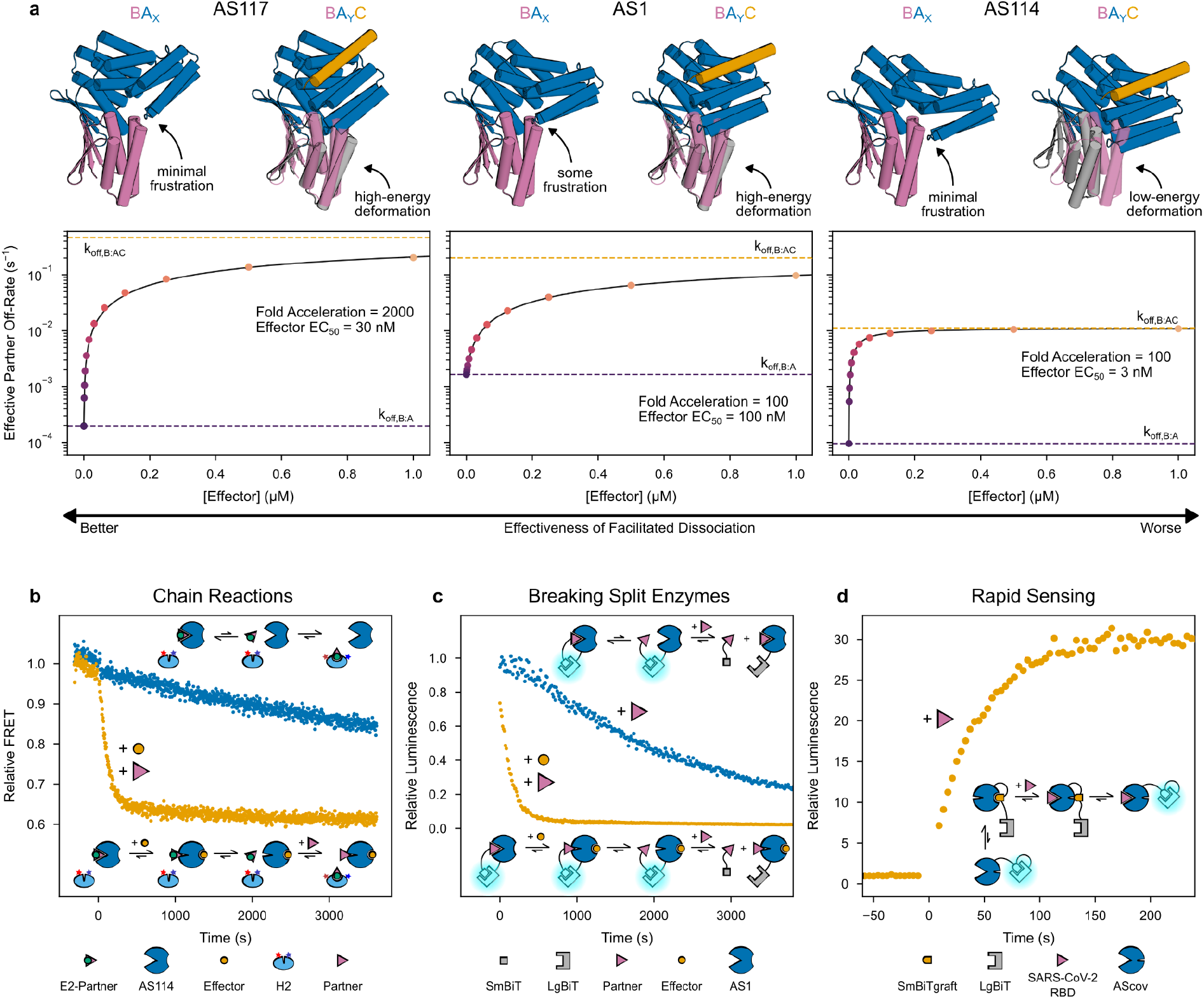
Modulation and applications of facilitated dissociation. **a**, Comparison of three representative designs with different facilitated dissociation behavior. (For each design) (top left) design model of host in state X (blue) aligned to the partner (pink) to show any clash influencing the partner off-rate in the absence of effector; (top right) design model of host-effector complex in state Y (blue and orange) aligned to the partner (pink) to show the designed allosteric clash, and (gray) AF2 prediction of the partner position relative to the switch in the ternary complex to show how the clash resolves through global strain; (bottom) effective partner off-rates plotted against effector concentration (circles) and fit with hyperbolic equations (black lines). **b**, Slow release of a kinetically trapped effector (upper schematic) and accelerated release through facilitated dissociation (lower schematic) with FRET time courses (normalized to the initial signal) of preincubated 500 nM AS114 and 250 nM E2-partner after adding 20 nM H2 then at time 0 adding 1 μM effector and excess partner (orange) or buffer (blue). A baseline drift (obtained from 500 nM AS114 after adding 20 nM H2 then at time 0 adding buffer) was subtracted from each time course. **c**, Breakage of a reversible split luciferase through slow direct competition (upper schematic) and faster facilitated dissociation (lower schematic) with luminescence time courses (normalized to the initial signal) of preincubated 100 pM AS1-LgBiT and 20 nM partner-SmBiT after adding 1 μM effector and 20 μM partner (orange) or just 20 μM partner (blue). **d**, Rapidly sensing SARS-CoV-2 through facilitated dissociation (schematic) with a luminescence time course (normalized to the initial signal) of 10 pM LgBiT-SmBiTgraft-AScov then at time 0 adding 800 nM SARS-CoV-2 RBD (orange).

We sought to model the structural basis for how strain in the ternary complex accelerates partner dissociation. The designs with the largest deformations in the strained ternary complex do not have the fastest dissociation rates: for example the predicted deformation magnitude for AS114 is double that of the fastest variant AS117 (Extended Data Fig. 4a,b), yet its accelerated partner dissociation is 40-fold slower (Fig. 4a). This could reflect the strain being so great in state Y that the ternary complex instead occupies state X, but DEER measurements of the AS1 and AS114 ternary complexes confirm that both primarily occupy state Y (Extended Data Fig. 2c). In a simple spring model of the ternary complex (Fig. 1d), lack of correlation between deformation magnitude and dissociation rates could arise if AS1 and the fast variants deform in directions of higher stiffness than the slower but more highly deforming variants like AS114. This anisotropy of stiffness may arise from the anisotropy of the secondary structure elements within the protein^32,33^: AF2 predicts AS1 and the fast variants to globally deform around axes nearly perpendicular to the binder interface helix but slower designs to deform around axes more parallel to this helix (Extended Data Fig. 4a,c). Using a simple coarse grained model based on the predicted magnitude and direction of the deformation, we find that across almost all designs, the predicted strain energy favoring partner dissociation correlates (R^2^ = 0.59) with the strain energy estimated from the observed rate enhancement (Extended Data Fig. 4d).

### Applications of Facilitated Dissociation Systems

We next set out to use facilitated dissociation to construct protein systems with kinetic behaviors previously inaccessible to design. First, inspired by dynamic DNA systems which function through toehold-mediated strand displacement^16^, we sought to create a kinetically-trapped system which, upon stimulation, quickly reconfigures through a chain reaction. To investigate this, we fused the effector peptide (E2) for a second hinge (H2) to the partner such that E2 is occluded when this fusion (E2-partner) is bound to a host (Extended Data Fig. 5). The release of E2-partner from AS114 and subsequent switching of H2 dramatically accelerates upon adding the original effector for AS114 and excess original partner (Fig. 4b, Extended Data Fig. 5). In principle, multiple orthogonal hosts could be constructed and chained together through these partner-effector fusions.

Second, we reasoned our designs could rapidly dissociate split protein systems that otherwise would be slow to reconfigure. To test this, we tagged AS1 and the partner with NanoBiT split luciferase fragments, LgBiT and SmBiT^34^. When combined, these components exhibit high luciferase activity which disappears much more rapidly upon addition of effector and excess untagged partner than upon addition of just excess untagged partner (Fig. 4c).

Third, in thermodynamically controlled biosensors designed previously, there is a tradeoff between dynamic range and response time primarily governed by a conformational pre-equilibrium^35,36^. We envisioned that our designs could form biosensors which directly couple the dynamic range to the analyte affinity without compromising response time. To investigate this, we grafted the SmBiT peptide onto the effector peptide at a range of positions, then screened these “SmBiTgraft” variants for binding to AS0 and rapid dissociation upon binding the partner. We used AS0 as the base design, reasoning that its closed state X would slow undesired peptide reassociation. For each working SmBiTgraft, we then generated a flexibly linked LgBiT-SmBiTgraft-AS0 construct. In these constructs, binding the partner should rapidly uncage the SmBiT to enable luciferase reconstitution. The best construct shows low luciferase activity which rapidly increases upon addition of partner, but only slowly upon addition of effector peptide, illustrating the kinetic advantage of facilitated dissociation over mutually exclusive competition (Extended Data Fig. 6). In principle, sensors for any analyte large enough to interact with sterically can now be constructed by swapping an analyte-specific binder into this platform. To demonstrate this, we rigidly fused a SARS-CoV-2 RBD binder (LCB1^37^) to the switch in place of the partner binder such that the RBD clashes with the switch in state Y but not in state X, and tested 16 of these “AScov” fusion designs. Upon addition of SARS-CoV-2 RBD, the best sensor shows a 30-fold increase in luciferase activity with a half-time of 30 seconds (Fig. 4d, Extended Data Fig. 6)—70 times faster than a previously designed LOCKR-based SARS-CoV-2 sensor which relies on mutually exclusive competition^35^. Thus, using this platform, a binder to almost any analyte can be turned into a single-component sensor that is sufficiently fast that in most practical applications, its response rate will be limited by analyte association rather than latch dissociation as in the LOCKR system.

### Rapid Modulation of IL-2 Signaling

Finally, we investigated whether facilitated dissociation could be used to control cellular processes with high temporal resolution. In cellular signaling, the residence time of the ligand stimulating the receptor is thought to modulate the signaling and cellular responses^38–40^. The central immune cytokine interleukin-2 (IL-2) activates the IL-2 receptor (IL-2Rβγ_c_) by inducing heterodimerization of chains β and γ_c_^41^. The resulting complexes dissociate on timescales of hours^42^, so controlling the temporal dynamics of IL-2 signaling is difficult: there is no off-switch (Fig. 5a).

**Fig. 5.**
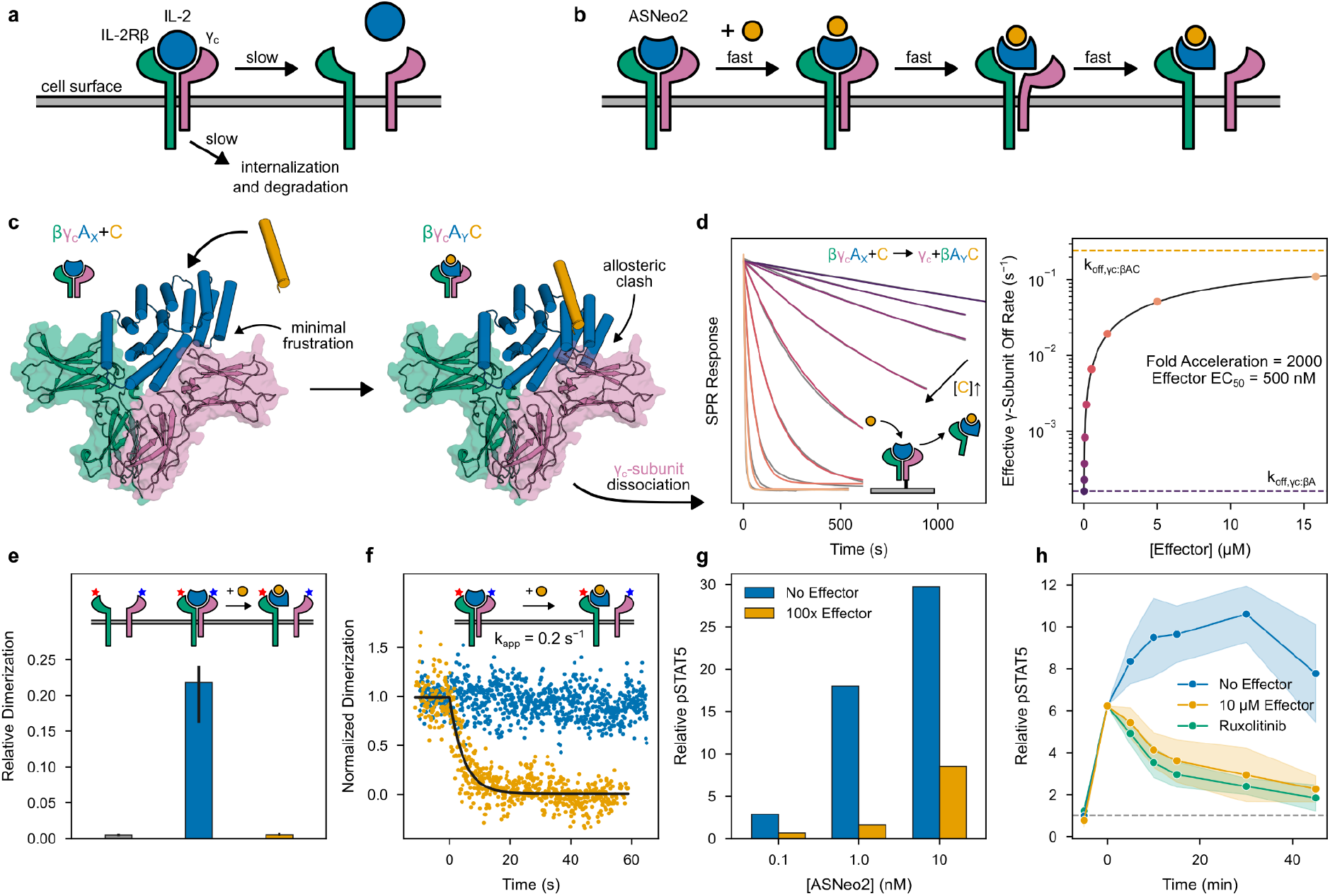
Design and characterization of a rapidly switchable IL-2 mimic. **a**, Natural pathways for switching off IL-2 signaling are slow. **b**, Design concept: through an induced-fit facilitated dissociation pathway, IL-2 signaling could be switched off rapidly. **c**, Design models of ASNeo2 in state X complexed with IL-2Rβγ_c_, the active signaling state (left), which can be quickly switched off by facilitated dissociation through a strained intermediate complex in state Y (right). **d**, (Left) fast effector concentration–dependent dissociation of γ_c_ upon addition of effector, data (gray) fit (colors) as described in methods. (Right) effective γ_c_ off-rates plotted against effector concentration (circles) and fit with a hyperbolic equation (black line). **e**, Median relative dimerization of IL-2Rβ and γ_c_ on the cell surface before adding ASNeo2 (gray), after adding 100 nM ASNeo2 (blue), and after later adding 10 μM effector (orange). Data combined from three independent experiments. Error bars represent 95% confidence intervals. Sample sizes are given in Extended Data Fig. 7. **f**, Time courses of dimerization of IL-2Rβ and γ_c_ after pre-stimulation with 100 nM ASNeo2 then adding nothing (blue) or 10 μM effector (orange) at room temperature. Each contains three independent time courses each normalized to its average initial relative dimerization. Data fit with a single exponential (black) yielding the rate constant *k*_app_. **g**, Dose-response of STAT5 phosphorylation relative to baseline upon stimulation with ASNeo2 (blue) or ASNeo2 precomplexed with 100x effector (orange) (one biological replicate). **h**, Time courses of STAT5 phosphorylation relative to baseline after stimulation with 1 nM ASNeo2 for 5 minutes then adding nothing (blue), 10 μM effector (orange), or 40 μM ruxolitinib (green) at 37°C. Three independent time courses were normalized to the signal at time 0, averaged, and renormalized to the average baseline signal. Shaded areas represent 95% confidence intervals. Some batches of cells were unresponsive to stimulation; data from these batches were excluded from analysis.

We set out to construct a switchable IL-2 mimic which enables seconds-timescale control over the formation of IL-2Rβγ_c_ (Fig. 5b). Neo-2, a previously designed IL-2 mimic, tightly binds IL-2Rβγ_c_ to elicit downstream signaling^43^. Sampling a variety of γ_c_ positions relative to the switch from AS1, we rigidly fused Neo-2 to the switch such that in state X, γ_c_ can bind, but in the effector-bound state Y, it would strongly clash (Fig. 5c). We identified several designs for which binding the effector dramatically accelerates dissociation of γ_c_ from the active signaling complex (Fig. S11). For the best of these, ASNeo2, binding the effector induces a 2000-fold increase in γ_c_ off-rate (Fig. 5d).

We used single-molecule fluorescence microscopy with labeled IL-2Rβ and γ_c_ to quantify receptor dimerization at the cell surface. Dual-color co-tracking and mobility analyses confirmed that ASNeo2 efficiently dimerizes IL-2Rβ and γ_c_, and that adding the effector rapidly and completely reverses this association even at elevated excess of γ_c_ (Fig. 5e,f; Extended Data Fig. 7c–i). By double-labeling ASNeo2 on opposite sides, we could observe the effector-induced intramolecular conformational change at the plasma membrane using single-molecule Förster resonance energy transfer (smFRET) (Extended Data Fig. 7j,k). ASNeo2 activates signaling in human NK (YT) cells, and its activity is greatly reduced in the presence of the effector (Fig. 5g). Upon adding the effector, STAT5 phosphorylation immediately stops accumulating and gradually decreases to a low level (Fig. 5h). The effector blocks ASNeo2 activity nearly as effectively as Ruxolitinib, a JAK1 inhibitor. The slightly lower potency of the effector could stem from minor activity of the effector-bound state or from a small population of cytokine which is unresponsive to the effector due to either partial degradation or internalization.

For downstream applications, it would be desirable to have a version that does not release active cytokine in the event of limited proteolysis. Splitting the Neo-2 across the switch would ensure that activity is lost if the switch regulating it degrades. ASNeo2 also requires higher effector concentrations than AS1 for a rapid response (Fig. 4a, 5d), suggesting the effector-binding cleft is predominantly collapsed in the βγ_c_-ASNeo2 complex. Using a split Neo-2 (Neo2A-Neo2B pair^44^) which minimally signals unless reconstituted, we repositioned the switch and Neo-2 to reduce strain in state βγ_c_A_X_, then rigidly fused Neo2B to the switch C-terminus and connected Neo2A to the switch N-terminus with a flexible linker (Extended Data Fig. 8). For several of these variants (especially ASNeo2_cp03), dissociation of γ_c_ from the active signaling complex dramatically accelerates under lower effector concentrations than required for the original design, suggesting that the effector-binding cleft is more intact (Extended Data Fig. 8). Dissociation from ASNeo2_cp08 is accelerated by 6000-fold, the highest fold change observed for any design here (Extended Data Fig. 8).

## Conclusion

We show that by explicitly considering excited intermediate states in designing protein structural transitions, protein-protein interaction kinetics can be controlled over a wide range of timescales. By using a flexible effector and a switch with an always-open effector-binding cleft, we created an induced-fit pathway for the effector to rapidly associate and mechanically stress a protein complex. By modulating the designed magnitude and direction of the resulting strain, we were able to control the accelerated dissociation kinetics of the binding partner. Crystal structures of our designs throughout the facilitated dissociation process demonstrate the accurate design of transiently populated states (the strained ternary complex) and large register-shift conformational changes. Our designs enable construction of kinetically governed systems (a previously challenging task), including chain reactions using energy stored in kinetic traps, split enzymes which associate tightly yet rapidly switch off, and sensors not limited by the usual tradeoff between response time and dynamic range. Each of these systems was successful on the first try without further optimization, indicating that given measurement of binding kinetics of the components alone, we can predict the behavior of larger composite systems.

Our kinetic data show that binding the peptide effector accelerates the switch conformational change by an induced-fit mechanism, whereas a more rigid effector can only bind via slow conformational selection. The peptide likely folds upon binding, effectively smoothing the energy landscape for the conformational transition: the energetic costs of uphill steps along the conformational change coordinate can be compensated by the formation of additional interactions with the folding peptide^30,45^. The kinesin power stroke also involves induced folding^46^, and our induced-fit conformational change has overall properties similar to power stroke mechanisms^8,26,27^, which can more efficiently generate force compared to ratchet mechanisms (in which a single large step is rectified at the end of the conformational change)^28^. In kinesin and other biological systems it is difficult to directly probe the role of effector flexibility and folding upon binding on overall function; our designed model systems in contrast allow direct comparison of flexible and rigid effectors and clearly demonstrate that the former yield considerably faster conformational transitions against loads.

Most known examples of facilitated dissociation couple the partner and effector through direct steric overlap^10,13,19,11,20,47,48^; in contrast, our approach amplifies rapid effector folding-upon-binding into a power stroke conformational change to allosterically accelerate partner dissociation. Because the allosteric coupling mechanism places no specific requirements on the partner, our approach can be used to dynamically regulate protein-protein interactions quite generally: by fusing to our switch, almost any binding interaction can be made to rapidly switch off in the presence of an effector. Our approach immediately transferred to rapidly switching active IL-2-like signaling complexes: we obtained multiple working designs on the first attempt when testing 24 designs. Transient IL-2 signaling with these switchable cytokines could enable investigation of early events in signaling, and since they switch off in response to a likely membrane-impermeable chemical stimulus, they could be used to decouple cell surface signaling from endosomal signaling to study differences in signaling from different cellular compartments. More generally, our work provides a route to designing not only protein structures, but also the rates and pathways of protein motion and change, which should ultimately enable construction of multistate protein machinery with complexity rivaling that powering life.

## Methods

### Design of structured switch-binder fusions (hosts) allosterically coupling the partner and effector

In PyMOL, we manually positioned the switch relative to the binder-partner complex subject to several constraints: there is no steric overlap between the partner and switch state X, there is large steric overlap between the partner and switch state Y, the smallest deformation that could be undergone by the switch and partner to resolve this clash is in the desired direction, and the switch C-terminus and binder N-terminus are relatively oriented such that sensible additional structure could be built between the switch and binder to rigidly fuse them as positioned. To aid in visualizing this additional structure, we also included placeholder helices while manually positioning the switch and binder, effectively “sketching” the fusion (Fig. S4a).

We then refined these sketches into actually plausible backbones. For the initial fusions including AS0, we extracted the center four residues of the placeholder helix, then used inpainting with RosettaFold^49^ to scaffold that fragment between the switch and binder. For later fusions including the AS1 variants and ASNeo2 designs, we first used Rosetta FastDesign^50^ to sample around the starting sketch for designable positions of the switch, binder, and placeholder helices while keeping the region of the state X switch that clashes in state Y fixed relative to the partner, then used RFDiffusion^7^ (conditioned on the secondary structure and block adjacency of the sketches) to build structure between the sampled switch and binder positions. During both structure generation approaches, we masked noncritical residues on the switch and binder interfacing with the fusion structure. Following generation of the fusion backbone structure, we used ProteinMPNN^51^ to optimize sequences for the fusion structure and the masked residues on the switch and binder. To filter designs, we used AF2^52^ (with initial guess (AF2-IG)^53^ for complex predictions) to predict the structure of the fusions alone, with the partner, and with the effector, selecting designs for which each structural state is correctly and confidently predicted by a majority of the five model weight sets. Finally, for the AS1 variants and ASNeo2 designs, we used AF2-IG to predict the structure of the strained ternary complexes, selecting sets with a diversity of deformation directions.

### Design of induced-fit register shift switches

Designs AS1, AS2, AS5, and AS7 were generated starting from design AS0, which contains the hinge cs221. When cs221 was designed, its state Y was generated by copying the N-terminal domain (helices 1–4) of its parent scaffold (DHR20^54^), aligning helix 4 of the copy to the corresponding helix of DHR20 offset by 3 residues, and combining the transformed N-terminal domain and original C-terminal domain into a single protein^24^. Thus, relative to the C-terminal domain, the N-terminal domain is both rotated around and translated along the axis of helix 4, exposing a cleft between the domains for binding a helical peptide. Now, to generate a new state X for this switch which retains an open cleft, we repeated this procedure but aligned the N-terminal domain with a residue offset of −4 instead of 3, so that overall this position of the N-terminal domain is shifted from state Y along helix 4 by one heptad (Fig. S5a). This introduces minimal rotation and thus maintains the open cleft. We combined this new position of the N-terminal domain with the entire C-terminal domain of AS0 (including the fusion to LHD101B), then used Rosetta FastDesign to further sample designable positions of the N-terminal domain around this starting point. In half the design trajectories, we included a placeholder helix in the cleft to help ensure it remains open. We then used inpainting with RosettaFold to generate loops connecting the domains, then paired these complete state X backbones with the original effector-bound state Y of AS0 (also generating new loops between domains in state Y as necessary to match the loop lengths of each new state X).

To generate single sequences supporting both states, we first performed multi-state design with Rosetta FastDesign (enforcing sequence symmetry between states) to further refine the paired backbones so they are more mutually compatible, then used ProteinMPNN with residue probabilities tied at corresponding positions between states to optimize sequences simultaneously for both conformations^24^. During these sequence design steps, the sequence of the effector and of the binder fusion was kept fixed. To filter designs, we used AF2-IG to predict the structure of the switches both with and without the effector, selecting designs for which the new state X (in the absence of effector) and state Y (in the presence of effector) are correctly and confidently predicted by a majority of the five model weight sets. To further ensure these designs favor state X in the absence of effector and state Y in the presence of effector, we only selected designs which scored more favorably in Rosetta in state X than in state Y, but also scored more favorably in state Y with the effector bound than the sum of the scores of state X and the unbound effector.

### Recombinant expression and purification

Synthetic DNA fragments encoding each design were obtained from IDT as eBlocks and cloned into custom vectors using Golden Gate assembly^55^. Designs usually carried a C-terminal sequence-specific nickel-assisted cleavage (SNAC) tag^56^ and 6xHis tag (MSG-Protein-GSGSHHWGSTHHHHHH). Proteins to be captured on the chip for SPR experiments carried an N-terminal AviTag™ and C-terminal 6xHis tag (MSGLNDIFEAQKIEWHESSG-Protein-GSGHHHHHH). For the SEC binding experiments shown in Fig. S6 and for screening SmBiTgraft effector peptide variants with SPR, the effector was fused to superfolder GFP in a sfGFP-GSSG-Effector-GSHHHHHH construct. To rapidly break split luciferases, AS1 was fused to LgBiT in a MSG-AS1-linker-LgBiT-GSHHHHHH construct and the partner was fused to SmBiT in a MSG-partner-linker-SmBiT-GSHHHHHH construct. In the rapid sensors, AS0 and AScov were fused to LgBiT and a SmBiT-containing effector in a MSGHHHHHHGS-LgBiT-linker-SmBiTgraft-linker-AS Protein-GS construct.

All proteins were expressed from NEB BL21(DE3) *E. coli* cells using TBII (MpBio) autoinduction media with 0.5% (w/v) glycerol, 0.05% (w/v) glucose, 0.2% (w/v) lactose, 20 mM MgSO_4_, trace metal mix, and 50 μg/mL kanamycin. 50 mL expression cultures were grown either at 37°C for 16–20 hr or at 37°C for 6–8 hr then at 18°C for 16–20 hours. Cells were harvested by centrifugation, resuspended in 5 mL lysis buffer (100 mM Tris HCl pH 8.0, 200 mM NaCl, 50 mM imidazole, 1 mM PMSF, 1 Pierce™ Protease Inhibitor Mini Tablets, EDTA-free per 50 mL), and lysed by sonication. The lysate was clarified by centrifugation at 14,000 x g for 30 min. Protein in the soluble lysate was bound to 1 mL Ni-NTA resin (Qiagen), washed with 5 mL low salt wash buffer (20 mM Tris HCl pH 8.0, 200 mM NaCl, 50 mM imidazole), 5 mL high salt wash buffer (20 mM Tris HCl pH 8.0, 1 M NaCl, 50 mM imidazole), and 5 mL low salt wash buffer, and eluted in 1.2 mL elution buffer (20 mM Tris HCl pH 8.0, 200 mM NaCl, 500 mM imidazole) after a 0.4 mL pre-elution. The proteins were further purified by SEC on an FPLC system with a Superdex 200 Increase 10/300 GL column in TBS (20 mM Tris pH 8.0, 100 mM NaCl) with 1 mL fractions. When possible, fractions probably corresponding to protein monomers were selected. Final protein concentrations were estimated using molar extinction coefficients predicted from the protein sequence and integrating the absorbance at 280 nm over the selected fractions. Correct protein molecular weights were confirmed using LC-MS.

### Peptide synthesis

The effector peptide cs221B was chemically synthesized by GenScript. The TAMRA-labeled effector used in FP experiments was synthesized as previously described^24^.

### Size-exclusion chromatography (SEC) binding assay

Individual host proteins, sfGFP-effector, and 1:1 host:sfGFP-effector mixtures were prepared at 20 μM in TBS (20 mM Tris pH 8.0, 100 mM NaCl). 0.5 mL of each solution was injected onto a Superdex 200 Increase 10/300 GL column in TBS and absorbance at 230 nm was monitored for changes in retention volumes of the mixture compared to the individual proteins.

### Fluorescence polarization (FP)

FP binding experiments with TAMRA-labeled effector were performed at 25°C in TBS (20 mM Tris pH 8.0, 100 mM NaCl) with 0.05% v/v TWEEN20 in 96-well plates (Corning 3686). Parallel and perpendicular fluorescence intensity was measured using a Synergy Neo2 plate reader with an FP 530/590 filter cube. Fluorescence polarization *P* (in units of mP) was calculated by the following expression.

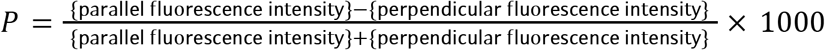

For affinity measurements, host proteins were titrated by two-fold serial dilution across TAMRA-labeled effector (at a constant concentration between 0.1 and 1 nM) through 24 wells, with a final volume of 80 μL in each well. Plates were incubated for at least 12 hours at room temperature to fully equilibrate before measurement. To determine affinities, the following binding isotherm function was fitted to the measured polarization values using nonlinear least-squares minimization.

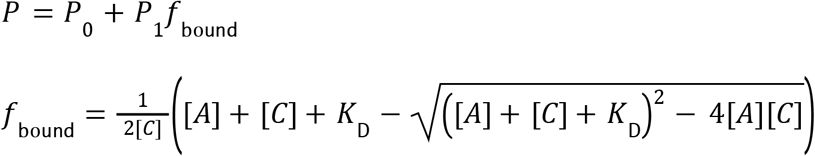

where *P* is the modeled polarization, *P*_0_ is the polarization of free effector, *P*_1_ is the change in polarization upon binding the host, *f*_bound_ is the fraction of effector bound to the host, [*A*] and [*C*] are the total concentrations of host and effector respectively, and *K*_D_ is the affinity between the host and effector. When the fit *K*_D_ is lower than [*C*], affinities are too strong to be accurately measured with this method, so *K*_D_ < [*C*] is reported.

For kinetic competition measurements, the partner LHD101An1 was titrated by two-fold serial dilution across the host protein (at a constant concentration of 22 nM) through 5 wells (a 6th well with just the host was also included) with a final volume of 72 μL in each well. Plates were incubated for 1 hour to allow the host and partner to equilibrate. To each well, 8 μL of 200 nM TAMRA-labeled effector was added and rapidly mixed using a multichannel pipette, and the measurement was started immediately afterward. This resulted in a 20 nM final concentration of both host and effector in 80 μL final volume per well. The following single exponential decay function was fitted to the measured polarization time courses using nonlinear least-squares minimization.

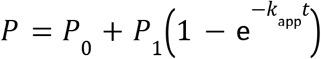

where *P* is the modeled polarization, *P*_0_ is the polarization of free effector, *P*_1_ is the amplitude of the change in polarization, *k*_app_ is the apparent rate constant, and *t* is the time after the start of the measurement.

### Surface plasmon resonance (SPR)

Proteins to be captured on the SPR chip were expressed with an N-terminal AviTag™ and purified as described above, except that the proteins were biotinylated following elution from the Ni-NTA resin: to the elutions, 5 μg/mL BirA (Avidity), 10 mM ATP, 10 mM Mg(OAc)_2_, and 100 μM d-biotin were added and allowed to incubate at room temperature for at least 4 hours prior to further purification by SEC. Successful biotinylation was confirmed using LC-MS. SPR measurements were performed at 25°C in HBS-EP+ buffer (Cytiva) on a Biacore 8K instrument. Biotinylated proteins were immobilized on the chip using the Biotin CAPture system (Cytiva).

For measurements of partner off-rates, biotinylated partner protein was immobilized on the chip. To measure base off-rates *k*_off,B:A_, host proteins at 50 nM were flowed over the chip for 60 s, then dissociation was measured for 2 hr. To measure accelerated off-rates *k*_off,B:AC_, preincubated host-effector complexes (host at 1 μM and effector at 5 μM to ensure the host is saturated with effector) were flowed for 60 s to form the ternary complex on the chip, then dissociation was measured for 4–20 min under constant flow of 5 μM effector. Most dissociation data was fitted with the following double exponential decay function to account for populations of host protein with different dissociation kinetics.

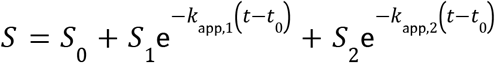

where *S* is the modeled SPR response, *S*_0_ is the baseline, *S*_1_ and *S*_2_ are amplitudes relating to the sizes of each host population, *k*_app,1_ and *k*_app,2_ are apparent rate constants corresponding to each host population, *t* is the time, and *t*_0_is the time at which dissociation initiates. The reported rate constant typically corresponds to the faster and higher amplitude exponential in the fit; instances where other criteria are used to determine which rate constant corresponds to the change of interest are noted. When clearly only one host population is present (often indicated by *k*_app,1_≈ *k*_app,2_ or a large difference between *S*_1_ and *S*_2_ when fitting a double exponential), the following single exponential decay function was fitted instead, where the parameters are the same as above. Instances where single exponentials were used are noted.

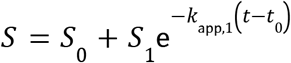

To measure the rate of effector association to form the ternary complex, biotinylated host protein was immobilized on the chip. The host protein was saturated with partner by flowing 5 μM partner over the chip for 4 min, then a varying concentration of effector and 5 μM partner was flowed over the chip for 2–4 min to associate the effector, and finally 5 μM partner was flowed over the chip for 4 min to monitor effector dissociation. Partner was included at 5 μM (higher than its affinity to the ternary complex) throughout the experiment to ensure the host remained saturated with partner, preventing changes in partner binding from convoluting the response from effector binding. Association data was fitted with the following single exponential decay function.

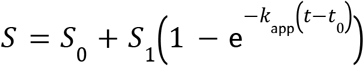

where *S* is the modeled SPR response, *S*_0_ is the baseline, *S*_1_ is the amplitude, *k*_app_ is the apparent rate constant, *t* is the time, and *t*_0_ is the time at which dissociation initiates.

With the peptide effector, the following linear function was fitted to the apparent rate constants.

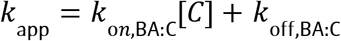

where *k*_*on*,BA:C_ and *k*_off,BA:C_ are on- and off-rate constants and [*C*] is the concentration of effector flowed over the chip.

With the 3hb effector, the following hyperbolic function was fitted to the apparent rate constants.

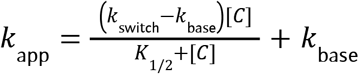

where *k*_switch_ is the rate constant for the BA_X_→BA_Y_ conformational change, *k*_base_ is a rate constant, and *K*_1/2_ is the concentration at which half the rate increase from *k*_base_ to *k*_switch_ is reached.

To measure the effector concentration-dependent rate of the full facilitated dissociation process, either biotinylated partner protein or biotinylated common gamma γ_c_ ectodomain (Acro Biosystems ILG-H85E8) was immobilized on the chip. For experiments with ASNeo2 designs, the ASNeo2 hosts were preincubated with IL-2Rβ ectodomain (Acro Biosystems CD2-H5221) before association with γ_c_. Each experiment involved multiple cycles of host association and induced dissociation under flow of various concentrations of effector obtained by two-fold serial dilution (Fig. S8a). Throughout these cycles, a small population of host that is unresponsive to the effector (“An”) could accumulate on the chip (Fig. S8a,b). The following system of differential equations describing the expected behavior of the proteins on the chip (accounting for this accumulation) can be fitted to the dissociation curve of cycle *n* (Fig. S8c).

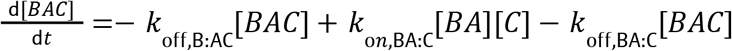

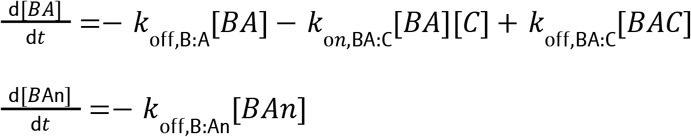

with initial values computed for each cycle by

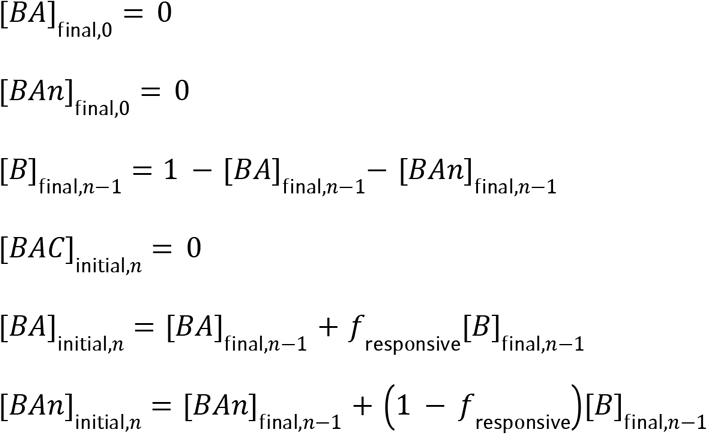

and the modeled concentrations of each complex state on the chip is related to the SPR response by

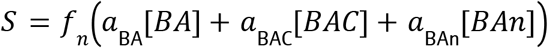

where *S* is the modeled SPR response, [*BAC*], [*BA*], and [*BAn*] are concentrations of complex states on the chip, *t* is the time after the start of the dissociation cycle, [*C*] is the concentration of effector flowed over the chip, *k*_off,B:A_, *k*_off,B:AC_, *k*_*on*,BA:C_, *k*_off,BA:C_, and *k*_off,B:An_ are on- and off-rate constants, *a*_BA_, *a*_BAC_, and *a*_BAn_ are amplitudes relating the concentration of each state on the chip to an SPR response, *f*_*n*_ is an amplitude fudge factor for cycle *n* accounting for small differences in amplitude across cycles, and *f*_responsive_is the fraction of host that is responsive to the effector. Varying all these parameters, this model is then globally fit to the dissociation curves of all cycles using nonlinear least-squares minimization. Note that with a sufficiently high value of *k*_*on*,BA:C_, this model’s dissociation kinetics are primarily determined by the partner dissociation parameters *k*_off,B:A_ and *k*_off,B:AC_; meanwhile, the effector binding parameters *k*_*on*,BA:C_ and *k*_off,BA:C_ tend to tightly covary and cannot be accurately determined from these fits. Also note that for designs with low values for *k*_off,BA:C_, this model may be less accurate because the assumption that [*BAC*] = 0 at the beginning of each cycle may no longer be valid. To compute the effective rate of the full facilitated dissociation process for intact host, we used the following simplified system of differential equations which no longer accounts for a small population of unresponsive host.

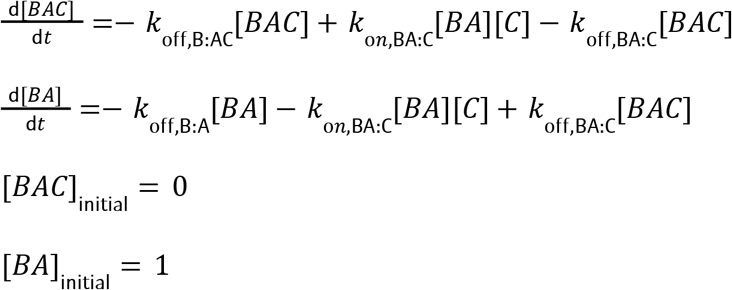

For each effector concentration [*C*], this system was solved for the half-time of the partner-host interaction *t*_1/2_ using the rate parameters determined from the original model fit to the data, and the effective rate of the full dissociation process *k*_eff_ was computed from each half-time as follows.

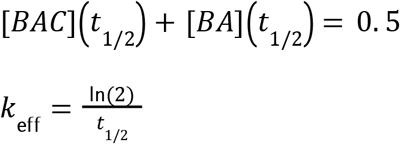

All SPR measurements in the main figures were repeated at least once, obtaining similar results.

### Circular dichroism (CD) spectroscopy

CD spectra were measured at 25°C on protein samples at 0.2 mg/mL in TBS (20 mM Tris pH 8.0, 100 mM NaCl) using a Jasco J-1500 spectrophotometer.

### X-ray crystallography

Protein was expressed from NEB BL21(DE3) *E. coli* cells using TBII autoinduction media as above but at a larger scale: either 8×50 mL or 1–2×500 mL cultures. Cells were harvested by centrifugation, resuspended in lysis buffer, and lysed by sonication. The lysate was clarified by centrifugation at 14,000 x g for 30 min. Protein in the soluble lysate was bound to 8 mL Ni-NTA resin (Qiagen), washed with 10 mL low salt wash buffer, 30 mL high salt wash buffer, and 10mL SNAC cleavage buffer (100 mM CHES, 100 mM Acetone oxime, 100 mM NaCl, 500 mM GnCl, pH 8.6)^56^, and incubated in 40mL SNAC cleavage buffer + 2 nM NiCl_2_ for 12 hours at room temperature to cleave. Afterwards, the flowthrough was collected and the beads were washed with 40 mL lysis buffer (minus the protease inhibitors). The amount of cleaved protein in the flowthrough and wash was assessed with SDS-PAGE, and fractions with enough cleaved protein were concentrated and further purified using SEC on an FPLC system with either a Superdex 75 Increase 10/300 GL column or a HiLoad 20/600 Superdex 75 pg column in either TBS (20 mM Tris pH 8.0, 100 mM NaCl) or lysine methylation buffer (50 mM HEPES pH 7.5, 250 mM NaCl) if lysines in the protein are to be methylated. For protein complex cocrystallization, the purified proteins and/or chemically synthesized effector were mixed at equimolar ratios. For some samples (resulting in crystals AS1_AB #1, AS1_AB #2, and AS1_ABC), lysines were methylated as previously described^57^, and the reaction was quenched using SEC on an FPLC system to buffer-exchange into TBS. Finally, the samples were concentrated to crystallization levels.

Crystallization experiments were conducted using the sitting drop vapor diffusion method. Initial crystallization trials were set up in 200 nL drops using the 96-well plate format at 20 °C. Crystallization plates were set up using a Mosquito LCP from SPT Labtech, then imaged using UVEX microscopes and UVEX PS-256 from JAN Scientific. Diffraction quality crystals formed in 0.2 M Magnesium chloride hexahydrate, 0.1 M Sodium cacodylate pH 6.5 and 50% v/v PEG 200 for AS1_A; in 0.2 M Sodium chloride, 0.1M Na/K phosphate pH 6.2 and 50% v/v PEG 200 for CS221B; in 0.1 M Sodium acetate pH 5.0, 5% w/v γ-PGA (Na+ from, LM) and 30% v/v PEG 400 for LHD101An1; in 0.2 M Magnesium chloride hexahydrate, 0.1 M Tris pH 8.5 and 25% w/v Polyethylene glycol 3,350 for AS5_AC; in 0.1 M Sodium acetate pH 5.0, 20% (v/v) MPD for AS5_A; in 0.2M 1,6-Hexanediol, 0.2M 1-Butanol, 0.2M 1,2-Propanediol, 0.2M 2-Propanol, 0.2M 1,4-Butanediol, 0.2M 1,3-Propanediol, Sodium HEPES; MOPS (acid) pH 7.5 and 40% v/v PEG 500* MME; 20% w/v PEG 20000 for AS1_AC; in 0.1 M Citric acid pH 3.5 and 2.0 M Ammonium sulfate for AS1_ABC; in 2.4 M Sodium malonate pH 7.0 for AS1_AB #1 (*P 6*_*1*_ *2 2*); and in 1.8 M Ammonium citrate tribasic pH 7.0 for AS1_AB #2 (*P 2*_*1*_ *2*_*1*_ *2*_*1*_).

Diffraction data was collected either at the National Synchrotron Light Source II on beamline 17-ID-1 (FMX/AMX) or Advanced Light Source beamline 821/822 or Advanced Photon Source NECAT 24ID-C. X-ray intensities and data reduction were evaluated and integrated using XDS^58^ and merged/scaled using Pointless/Aimless in the CCP4 program suite^59^. Structure determination and refinement starting phases were obtained by molecular replacement using Phaser^60^ using the designed model for the structures. Following molecular replacement, the models were improved using phenix.autobuild; with rebuild-in-place to false, and using simulated annealing. Structures were refined in Phenix^61^. Model building was performed using COOT^62^. The final model was evaluated using MolProbity^63^. Data collection and refinement statistics are recorded in Supplementary Table 3.

### Double electron-electron resonance (DEER) spectroscopy

Spin label modeling and distance distribution predictions were performed as previously described^24^ using chiLife^64^ with the off-rotamer sampling method^65^. Site pair selections were performed as previously described^24^. Host protein variants containing cysteines at the selected sites were purified as described above, except that 0.5 mM TCEP was included in the lysis buffer and the first two Ni-NTA resin washes, and the proteins were labeled immediately after elution from the Ni-NTA resin: to the elutions, 50 μL of 200 mM MTSL in DMSO was added and allowed to incubate for at least 2 hours at room temperature prior to further purification by SEC. Successful labeling was confirmed using LC-MS. DEER samples were prepared at 20 μM labeled host protein in deuterated solvent buffered by 20 mM Tris at pH 8.0 with 100 mM NaCl and 20% d_8_-glycerol (Cambridge Isotope Laboratories, Inc.) as a glassing agent. When appropriate, partner and effector were added to a concentration of 100 μM each. 15-30 μL of each sample were loaded into a quartz capillary (Sutter Instrument, 1.1 mm inner diameter, 1.5 mm outer diameter) and flash frozen with liquid nitrogen. Samples were stored at −80°C until the DEER experiment was conducted.

DEER experiments were performed as previously described^24^. An ELEXSYS E580 spectrometer (Bruker) at Q-band (∼34 GHz) with an EN5107D2 resonator (Bruker) was used for all experiments. Temperature was maintained at 50 K using a cryogen-free cooling system (ColdEdge). The 4-pulse DEER sequence was used using 60 ns Gaussian observer pulses with a full width at half maximum (FWHM) of 30 ns and a frequency near the center of the field-swept spectrum and 150 ns sech/tanh probe pulses with center 80 MHz above the observer frequency, 80 MHz bandwidth, and a truncation parameter of 10. All shaped pulses were generated using the SpinJet arbitrary waveform generator (Bruker). Pulse shapes were calculated using PulseShape (https://github.com/stolllab/PulseShape) using both resonator compensation and transmitter nonlinearity compensation. All data were collected using a 2 ms shot repetition time, 8-step phase cycling, and *τ*_1_ averaging from 400 ns to 528 ns in 16 ns steps. All additional experimental parameters, including pump pulse time step, *τ*_2_ delays, number of scans, and more were chosen on a per-sample basis and are reported in Supplementary Table 2.

All DEER data were analyzed using the DeerExp module of the eprTools Python package (https://github.com/mtessmer/eprTools). All data were fit using separable non-linear least squares^66^. The foreground signal was modeled using Tikhonov regularization with the second derivative operator. The regularization parameter was selected using generalized cross-validation. The background was modeled using a 3D-homogeneous spin distribution. An additional penalty restraining the modulation depth to be low was used to prevent the fitting of long-distance artifacts in the foreground as previously done^67^. Confidence intervals were estimated using bootstrap sampling with 100 samples using a fixed regularization parameter. Fit parameters such as the regularization parameter, the modulation depth, and the signal-to-noise ratio are listed in Supplementary Table 2.

### Molecular dynamics (MD) simulations

Input files for the MD simulation were prepared with CHARMM-GUI^68,69^ with AlphaFold2 structure as the initial structure. The rectangular water box of edge length 10Å was placed around the protein. The water box contained potassium and chlorine ions of concentration 0.15M that neutralized the protein’s net charge. The ions were placed in the water box using Monte Carlo method. To model the system, the CHARMM36m force field^70^ was used. After the explicit solvent system was made, the system was minimized and equilibrated before the production MD run. All three steps were performed with the GROMACS 2020.2 MD engine^71,72^.

Steepest descent method was used for energy minimization, for 5000 steps with an energy tolerance of 1000 kJ/mol/nm. Neighbor list was updated every 10 steps with a cut-off distance of 1.2 nm. Cut-off was used to calculate Van der Waals interactions with switch distance of 1.0 nm and cut-off distance of 1.2 nm. The force was smoothly switched off between the switch distance and the cut-off distance. The fast smooth particle-mesh Ewald method was used to calculate electrostatics with a cut-off distance of 1.0 nm. All bonds with hydrogen atoms were treated as rigid using the Linear Constraint Solver (LINCS) algorithm^73^.

The equilibration step was performed with a leap-frog algorithm using a timestep of 1 fs and total simulation time of 125 ps. The system was propagated in the NVT ensemble. The temperature was maintained at 303.15K using the velocity rescaling method. The solute and solvent were coupled with a time constant for coupling of 1 ps. The center of mass translational velocity was removed every 100 steps to prevent the system from drifting. The same cut-off scheme for Van der Waals and electrostatic interactions were used as in the minimization step except the neighbor list was updated every 20 steps. The velocities were generated from Maxwell distribution with temperature of 303.15K. The same LINCS method was used to constrain the hydrogen atoms during the equilibration step.

The production step was performed with a leap-frog algorithm using a timestep of 2 fs and total simulation time of 1 μs. The trajectories with the same initial equilibrated structure were obtained in triplicate. The system was propagated in the NPT ensemble. Temperature was maintained at 303.15K using velocity rescaling method. The solute and solvent were coupled with a time constant of 1 ps. Exponential relaxation pressure coupling and isotropic coupling with time constant of 5.0 ps was used to maintain pressure at 1.0 bar. The same cut-off scheme for Van der Waals and electrostatic interactions were used as in the minimization step except the neighbor list was updated every 20 steps and the Coulomb cut-off distance was set as 1.2 nm. The same center of mass velocity removal was used in the production step as in the equilibration step. The same LINCS method was used to constrain the hydrogen atoms during the production step.

Post analysis of the trajectories such as RMSD and RMSF calculation was performed with MDAnalysis^74^. To simulate DEER distance distributions from these trajectories, structures along each trajectory were clustered using the Gromos clustering algorithm^75^ in GROMACS, distance distributions were predicted (using chiLife^64^ with the off-rotamer sampling method^65^) for the center structure of each cluster, and the resulting distributions were averaged weighted by the occupancies of their corresponding clusters.

### Chain reactions with FRET readout

Hinge protein cs201F_E249L (H2) was purified as described above, except that 0.5 mM TCEP was included in the lysis, wash, and elution buffers, and the SEC buffer was PBS (20 mM sodium phosphate pH 7.0, 100 mM NaCl, 0.5 mM TCEP). To double label with dyes, 50 μM H2 was incubated with 250 μM Alexa Fluor™ 555 C_2_ Maleimide (donor, Thermo Fisher Scientific) and 250 μM Alexa Fluor™ 647 C_2_ Maleimide (acceptor, Thermo Fisher Scientific) shaking at room temperature for at least 2 hours. The reaction was quenched by adding DTT to 10 mM, and the proteins were separated from excess dye by SEC in TBS (20 mM Tris pH 8.0, 100 mM NaCl).

FRET binding experiments were performed at 25°C in TBS (20 mM Tris pH 8.0, 100 mM NaCl) with 0.05% v/v TWEEN20 in 96-well plates (Corning 3686). Fluorescence intensity was measured using a Synergy Neo2 plate reader, exciting the donor at 520 nm wavelength and reading acceptor emission at 665 nm wavelength.

To measure E2-partner on-rate, 40 μL H2 at 10 nM was prepared in 6 wells. To each well, 40 μL E2-partner at various concentrations was added and rapidly mixed using a multichannel pipette, and the measurement was started immediately afterward. The following single exponential decay function was fitted to the measured FRET time courses using nonlinear least-squares minimization.

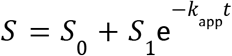

where *S* is the modeled fluorescence signal, *S*_0_ is the fluorescence at equilibrium, *S*_1_ is the amplitude of the change in fluorescence, *k*_app_ is the apparent rate constant, and *t* is the time after the start of the measurement. The following linear function was fitted to the apparent rate constants.

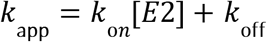

where *k*_on_ and *k*_off_ are on- and off-rate constants and [*E*2] is the total concentration of E2-partner.

To measure the accelerated transfer of E2 from AS114 to H2, 1067 nM AS114 and 533 nM E2-partner were incubated for 15 minutes to fully cage E2-partner in AS114. A control solution of just 1067 nM AS114 was also prepared. 37.5 μL of 42.7 nM H2 was prepared in 8 wells. 37.5 μL of AS114+E2-partner was added to 4 wells and 37.5 μL of AS114 was added to the other 4 wells using a multichannel pipette, and the measurement was started immediately afterward. 5 min later, 5 μL of buffer, 100 μM partner, 16 μM effector, or 100 μM partner and 16 μM effector were added to each set of 4 wells using a multichannel pipette, and the measurement was immediately continued for 1 hr. Mixing components at these concentrations resulted in final concentrations of 500 nM AS114, 250 nM E2-partner, 20 nM H2, 1 μM effector, and 6 μM partner. A baseline drift function of the following form was fit to the AS114+buffer data and subtracted from the other time courses.

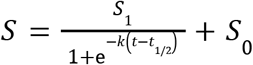

where *S* is the modeled fluorescence signal, *S*_0_ is the initial fluorescence, *S*_1_ is the amplitude of the change in fluorescence, *k* is a rate constant, *t* is the time after the start of the measurement, and *t*_1/2_ is the time at which the fluorescence has changed by half the full amplitude.

### Rapid sensors and split enzymes with luminescence readout

Luminescence experiments were performed at 25°C in TBS (20 mM Tris pH 8.0, 100 mM NaCl) with 0.05% v/v TWEEN20 in 96-well plates (Corning 3686). Luminescence was measured using a Synergy Neo2 plate reader with a LUM filter cube.

To measure rapid breakage of a split luciferase, 111 pM AS1-LgBiT and 22 nM partner-SmBiT were incubated for 1 hr to load AS1 with the partner and reconstitute the split luciferase. 72 μL of this mixture was added to 2 wells. 8 μL of 1/10 diluted Nano-Glo® substrate (Promega N1130) and either 10 μM effector and 200 μM partner or just 200 μM partner was added to these wells using a multichannel pipette, and the measurement was started immediately afterward. Mixing components at these concentrations resulted in final concentrations of 100 pM AS1-LgBiT, 20 nM partner-SmBiT, 1 μM effector, 20 μM partner, and 1/100 diluted Nano-Glo® substrate.

To measure analyte sensing, 64 μL of 12.5 pM sensor was added to 8 wells. 8 μL of 1/10 diluted Nano-Glo® substrate was added to these wells using a multichannel pipette, and the measurement was started immediately afterward. 5 min later, 8 μL of various concentrations of either partner or effector obtained by ten-fold serial dilution were added using a multichannel pipette, and the measurement was immediately continued for 30–60 min. Mixing components at these concentrations resulted in final concentrations of 10 pM sensor and 1/100 diluted Nano-Glo® substrate.

### Live cell single molecule imaging

For cell surface labeling, receptors were N-terminally fused to suitable tags using a pSems vector including the signal sequence of Igκ (pSems-leader). Common gamma chain (γc) was fused to the ALFA-tag^76^ and IL-2Rβ was fused to nonfluorescent monomeric GFP (mXFP)^77^. HeLa cells (ACC 57, DSMZ Germany) were cultured as previously described^78^. For transient transfection, cells were incubated for 4–6 h with a mixture of 150 mM NaCl, 10 μL of 1 mg/mL polyethylenimine (PEI MAX®, Polysciences 24765) and 200ng of DNA of pSems leader ALFAtag-γc and 2800ng of pSems leader mXFPe1-IL-2Rβ^79^. Labeling, washing and subsequent imaging were performed after mounting the coverslips into custom-made incubation chambers with a volume of 1 ml. Cells were equilibrated in medium with FBS but lacking phenol red supplemented with an oxygen scavenger and a redox-active photoprotectant (0.5 mg mL^−1^ glucose oxidase (Sigma-Aldrich), 0.04 mg mL^−1^ catalase (Roche), 5% w/v glucose, 1 μM ascorbic acid and 1 μM methylviologene) to minimize photobleaching^80^.

Selective cell surface receptor labeling was achieved by using anti-GFP and anti-ALFAtag Nanobodies (NBs), which were site-specifically labeled by maleimide chemistry via a single cysteine residue at their C-termini^80^. Anti-ALFA NB labeled with Cy3B (DOL; 1.06) and anti-GFP NB labeled with ATTO 643 (DOL; 1.0) were added at concentrations of 3 nM each, at least 10 min before imaging. Coverslips were precoated with poly-l-lysine-graft-poly(ethylene glycol) to minimize unspecific binding of NBs and functionalized with RGD peptide for efficient cell adhesion^81^.

Dual-color imaging was carried out by total internal reflection fluorescence microscopy using an inverted microscope (IX-83, Olympus) equipped with a spectral image splitter (QuadView, Photometrics) and an EMCCD camera (iXon Ultra, Andor) as described in detail elsewhere^82^. Fluorophores were excited by sequential illumination with a 561-nm laser (2RU-VFL-P-2000-560-B1R, 2000W, MPB communications) and a 642-nm laser (2RU-VFL-P-2000-642-B1R, 2000W, MPB communications). Alternating laser excitation was achieved with a simple micro-controller (Arduino Uno) and open-source acquisition software^83^ synchronizing laser shuttering via an AOTF (AA.AOTFnC-400.650-TN, AA Opto Electronic) and camera triggering. For long term tracking experiments, 1500 frames per channel were acquired at 40 fps. Resulting image stacks were divided into 5-frame stacks and dimerization was determined for each stack. For all other tracking experiments, 200 frames per channel were acquired at 40 fps and dimerization was determined over the whole image stack.

Dual-color single molecule co-tracking time-lapse images were evaluated using an in-house developed MATLAB software (SLIMfast4C, https://zenodo.org/record/5712332)^80^. After channel registration based on calibration with fiducial markers, molecules were localized using the multi-target tracking algorithm^84^. Immobile emitters were filtered out by spatiotemporal cluster analysis^85^. Frame-by-frame co-localization within a cut-off radius of 150 nm was applied followed by tracking of co-localized emitters using the utrack algorithm^86^. Molecules co-diffusing for 10 frames or more were then identified as co-localized. Relative levels of co-localization were determined based on the fraction of co-localized particles relative to all localizations in the ALFA-γc channel (561 nm). Diffusion properties were determined from pooled single trajectory using mean squared displacement analysis for all trajectories with a lifetime greater than 10 frames. Diffusion constants were determined from the mean squared displacement by linear regression. Relative dimerization was estimated by:

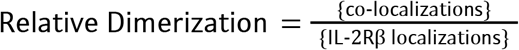

FRET efficiencies were evaluated using an in-house developed MATLAB software (provided and described in detail elsewhere as supplementary software^82^). In brief, alternating laser excitation FRET experiments provide three separated emission channels: directly excited donor 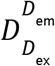 and acceptor 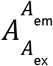 as well as a sensitized FRET 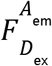 channels. First, channels were aligned based on calibration with fiducial markers. Then, after applying a single molecule localization algorithm^87^, single-molecule intensities were determined from background subtracted images 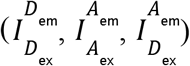. To evaluate FRET efficiencies, donor-acceptor pairs were co-localized based on an optimized search radius. For these pairs, the apparent FRET Efficiency was calculated by:

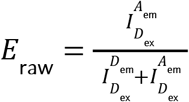

In order to achieve accurate FRET efficiencies, standard further corrections were applied. These include the donor leakage coefficient, crosstalk corrected proximity ratio and the correction factor γ^82,88^.

### YT signaling assay

YT-1 cells were cultured in RPMI 1640 complete medium, supplemented with 10% fetal bovine serum, 2 mM L-glutamine, minimum essential non-essential amino acids, sodium pyruvate, 25 mM HEPES, and penicillin-streptomycin (Gibco). For the flow cytometry-based pSTAT5 detection assay, 2–5 × 10^5^ IL-2Rα positive YT-1 cells were seeded in 350 μL of medium per well of a 96-well plate. The cells were stimulated with 1 nM ASNeo2 or Neo2 for 5 minutes at 37°C. As a control, 50 μL of untreated YT-1 cells were set aside at the start of each experiment and evaluated after 45 minutes alongside the treated cells. After stimulation, all cells were transferred to three separate wells containing either a control (no treatment), 10 μM Effector or 40 μM Ruxolitinib. One-seventh of the cells (i.e., 50 μl) were resuspended in 17 μl of 16% paraformaldehyde (PFA) for immediate fixation. This process was repeated at 5, 10, 15, 30, and 45-minute intervals. After all the time points were fixed, the cells were re-fixed in 4% PFA for 15 minutes at room temperature. Following fixation, the cells were washed once with PBS containing 0.5% BSA (PBSA) and permeabilized with 100% methanol for 45 minutes at 4°C. After permeabilization, the cells were washed twice with PBSA and stained for 1 hour at room temperature using Alexa Fluor® 647-conjugated Phospho-STAT5 (Tyr694) rabbit monoclonal antibody (Cell Signaling Technology, #9365). After three washing steps, the cells were analyzed using a CytoFlex S flow cytometer (Beckman Coulter). Data were analyzed with CytExpert software, and cells were gated based on SSC-A vs FSC-A. Each experiment was conducted in triplicate, and the results were analyzed accordingly.

## Acknowledgements

We thank P. J. Y. Leung, K. L. Shelley, A. Pillai, C. Demakis, M. Exposit, R. J. Ragotte, and M. Glögl for helpful discussions and technical support, K. VanWormer and L. Goldschmidt for technical support, S. R. Gerben and A. Murray for protein production support, and X. Li, M. Lamb, Z. Taylor, and V. Adebomi for LC-MS support. This work was supported by the Audacious Project at the Institute for Protein Design (A.J.B., A.K., J.D.L.C., E.B., and A.K.B.), by a gift from Microsoft (A.J.B.), by the Nordstrom Barrier Institute for Protein Design Directors Fund (M.H.A. and F.P.), by Bill and Melinda Gates Foundation #OPP1156262 (A.K., J.D.L.C.), by the Open Philanthropy Project Improving Protein Design Fund (E.B., A.K.B.), by the National Institutes of Health’s National Institute of Allergy and Infectious Disease grant R0AI160052 (A.K.B.), by National Science Foundation grant MCB 2119837 and National Institutes of Health grant GM115805 (W.H.R. and D.M.Z.), by National Institutes of Health grant GM151956 (S.S.), by the DFG grants PI 405/15 and SFB 1557 (C.P. and J.P.), and by the Howard Hughes Medical Institute (A.K.B. and D.B.). The EPR spectrometer used for the DEER experiments was in part supported by National Institutes of Health grant S10OD021557. This research used resources (FMX/AMX) of the National Synchrotron Light Source II, a U.S. Department of Energy (DOE) Office of Science User Facility operated for the DOE Office of Science by Brookhaven National Laboratory under Contract No. DE-SC0012704. The Center for BioMolecular Structure (CBMS) is primarily supported by the National Institutes of Health, National Institute of General Medical Sciences (NIGMS) through a Center Core P30 Grant (P30GM133893), and by the DOE Office of Biological and Environmental Research (KP1607011). This work is based upon research conducted at the Northeastern Collaborative Access Team beamlines, which are funded by the National Institute of General Medical Sciences from the National Institutes of Health (P30 GM124165). This research used resources of the Advanced Photon Source, a U.S. Department of Energy (DOE) Office of Science User Facility operated for the DOE Office of Science by Argonne National Laboratory under Contract No. DE-AC02-06CH11357. The Berkeley Center for Structural Biology is supported by the NIH, National Institute of General Medical Sciences, and the HHMI. The ALS is supported by the Director, Office of Science, Office of Basic Energy Sciences and US Department of Energy (DOE) (DE-AC02-05CH11231).

## Competing Interests

The authors declare the following competing interests: A.J.B., F.P., A.K.B., and D.B. are in the process of filing a provisional patent application that incorporates discoveries described in this article.

## Author Contributions

A.J.B. developed the facilitated dissociation design concept, developed the computational design pipeline, and designed and characterized the allosteric switches. C.P. performed and analyzed single molecule localization microscopy and final signaling activity experiments. M.A.L. and A.J.B. designed and characterized the rapid sensors. M.D.J. and M.H.T. performed and analyzed DEER experiments. W.H.R and A.J.B. performed and analyzed MD simulations. A.J.B. and M.H.A. performed initial signaling activity experiments. D.D.S. and A.A. performed site-saturation mutagenesis on LHD101. A.K., J.D.L.C., E.B., B.S., and A.K.B. determined crystal structures. D.B., J.P., F.P., S.S., and D.M.Z. supervised research. A.J.B. and D.B. wrote the manuscript. F.P., C.P., M.A.L., M.H.T., and W.H.R. contributed to the manuscript. All authors read and commented on the manuscript.

**Extended Data Fig. 1:**
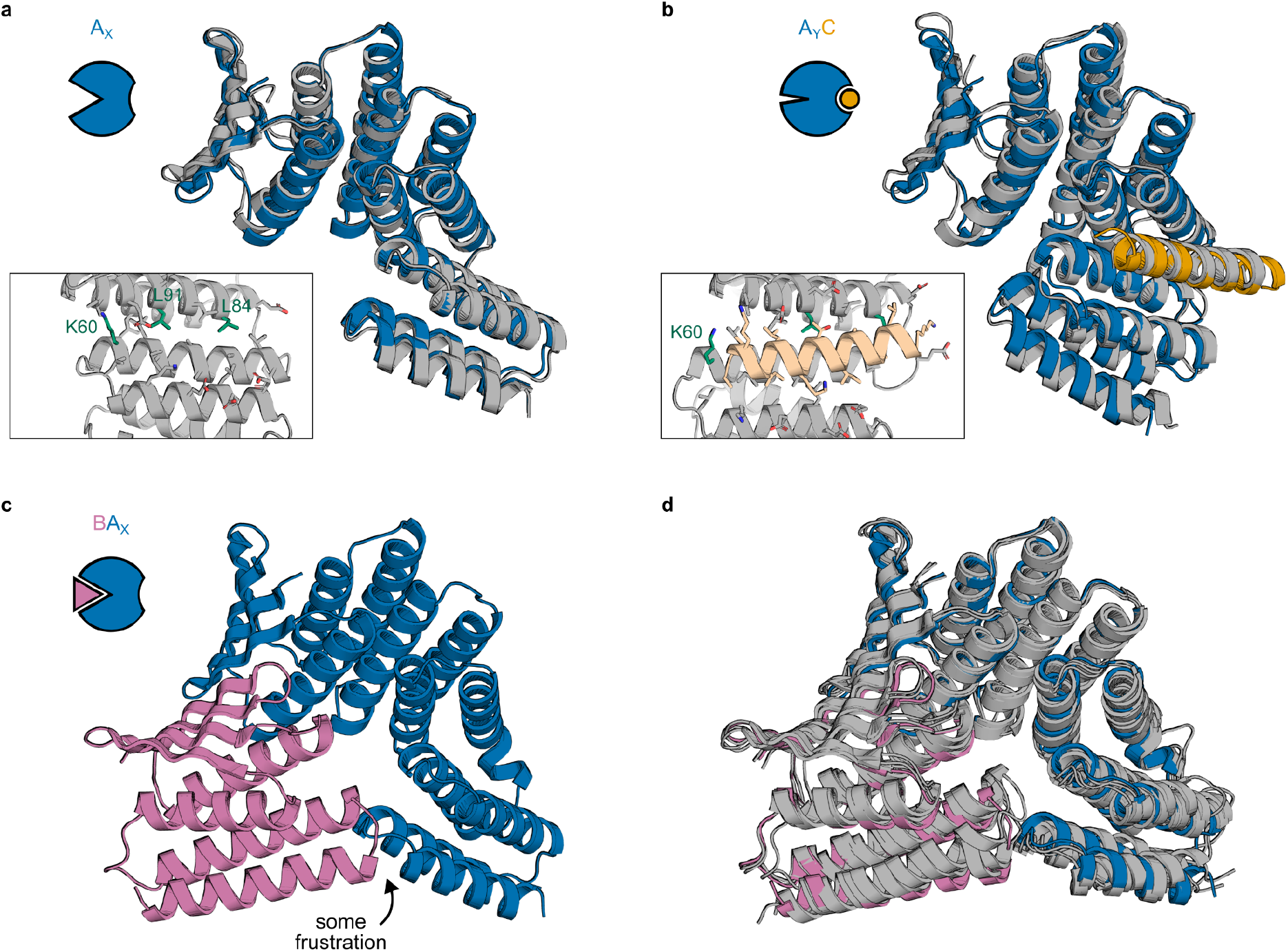
Structural characterization of AS5 and structural frustration of AS1 state BA_X_. **a**, Crystal structure of AS5 alone (gray) overlaid with the design model of AS5 in state X (blue). Inset shows a detailed view of side chains in the partially-open effector-binding cleft. **b**, Cocrystal structure of AS5 and peptide effector (gray) overlaid with the design model of the AS5-effector complex in state Y (AS5 in blue, effector in orange). Inset shows the same view of the side chains in the effector-binding cleft as in (a). **c**, Design model of AS1 in state X (blue) aligned to the partner (pink), showing a minor clash. **d**, Three cocrystal structures of AS1 (with intact cleft) and partner with methylated lysines (gray) overlaid with the AF2 model of the partner-AS1 complex in state X (AS1 in blue, partner in pink), showing fluctuation in the partner binding conformation.

**Extended Data Fig. 2:**
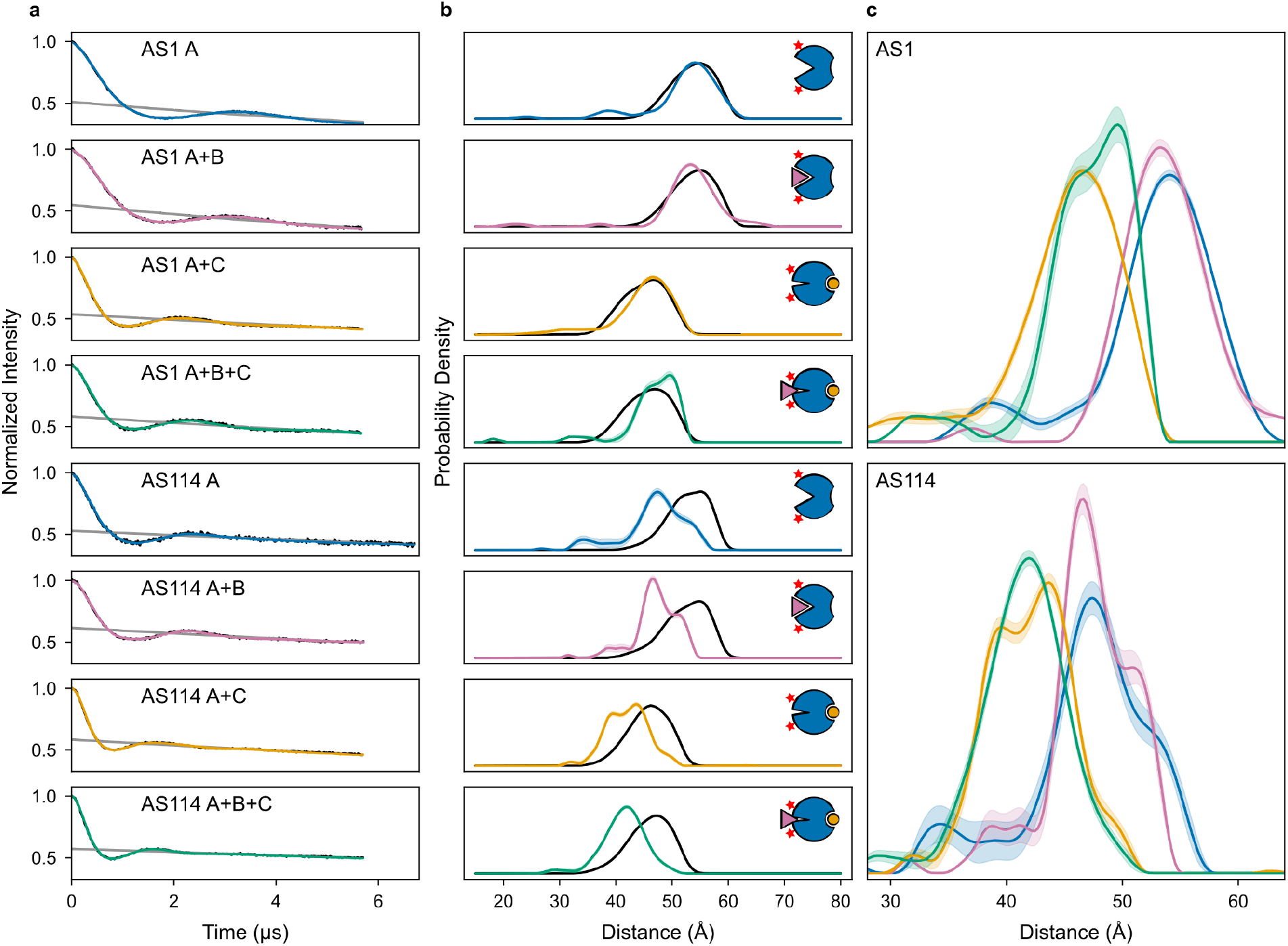
DEER characterization of AS1 and AS114. **a**, Raw DEER traces (black), foreground fits (colors), and background fits (gray) for AS1 and AS114 with all combinations of partner and effector. Experiments on complexes included the partner, effector, or both in excess over the host at concentrations higher than required to fully form the complex (Fig. S9), and spin labels were placed far from the partner and effector binding sites. Thus, changes in the DEER distance distributions with different combinations of partner and effector should only reflect changes in host conformation. **b**, Distance distributions from experiment (colors with shaded 95% confidence intervals) and simulated from the structural state represented by the cartoons (black). For AS1, the simulated and experimental distance distributions agree well, further validating that each state adopts its designed conformation. For AS114, the simulations consistently overestimate the experimental distribution by ∼5 Å, but the shift in the distance distributions with the effector compared to those without validates the designed conformational change. **c**, Experimental distance distributions of all states, colored corresponding to (b) and with shaded 95% confidence intervals. For both AS1 and AS114, the ternary complex distribution (green) aligns with the host-effector complex distribution (orange) and not with the host alone (blue) or partner-host complex (pink) distributions, confirming that the ternary complex is primarily in state Y.

**Extended Data Fig. 3:**
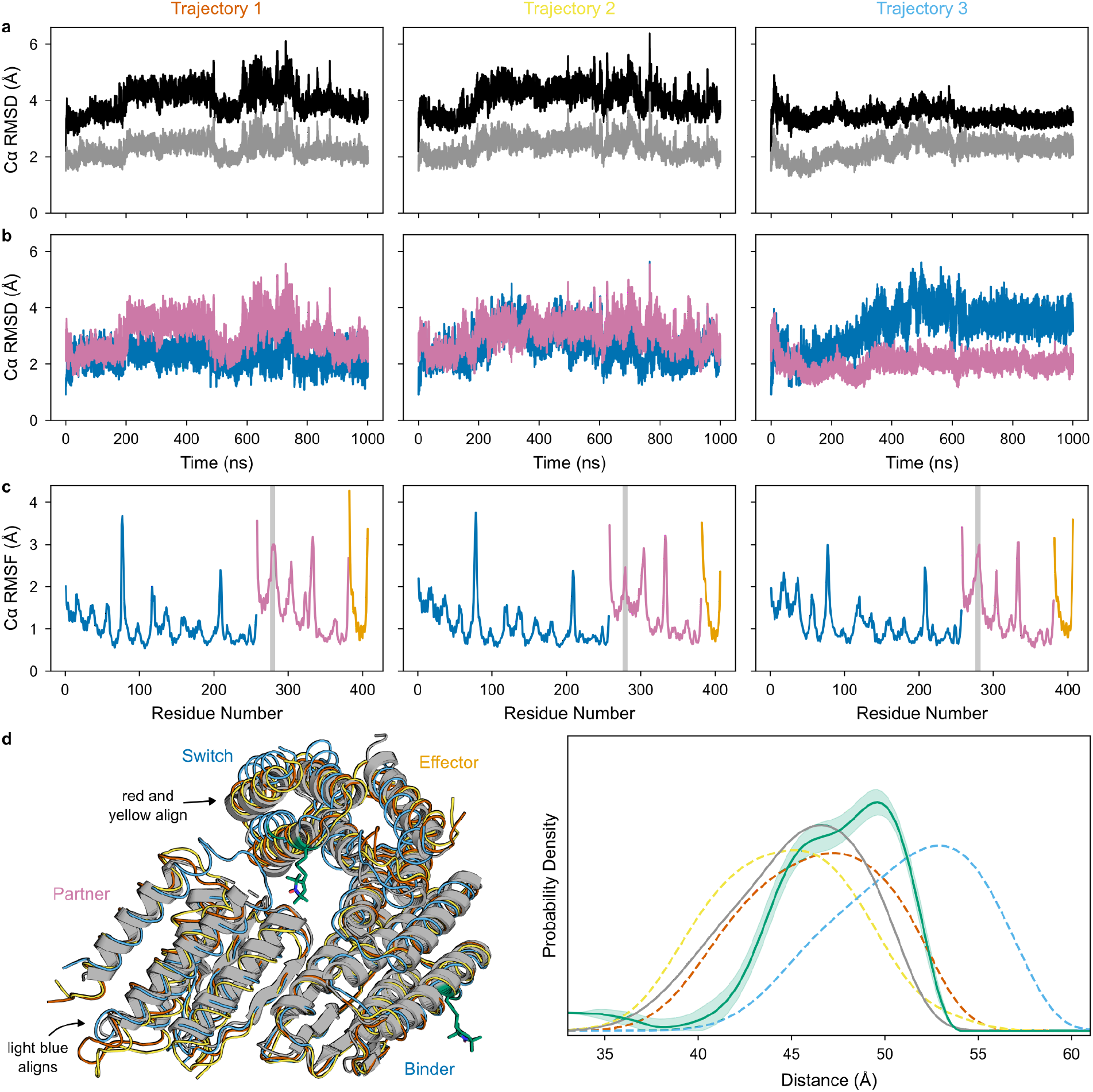
MD characterization of the AS1 ternary complex. **a**, Lower Cα RMSD of the MD trajectories from the crystal structure (gray) than from the aligned clashing design models of partner and host-effector complex (black), showing that the MD simulations strain away from the clashing state in a manner similar to the crystal structure. **b**, Cα RMSD of the switch (blue) and partner (pink) from their position in the crystal structure when the entire structures are aligned. Compared to trajectories 1 and 2, trajectory 3 shows reduced partner deformation and increased switch deformation, showing that these trajectories differ in where they localize strain to resolve the clash. **c**, Per-residue Cα RMSF of the host (blue), partner (pink), and effector (orange) in the ternary complex computed from each trajectory. The clashing region of the partner (highlighted in gray) shows significant flexibility, according with this region being disordered in the crystal structure. **d**, Comparison of MD simulations to experimental data. (Left) Crystal conformation of the ternary complex (gray) aligned to representative conformations from each MD trajectory (red, yellow, and light blue). DEER spin label positions are shown in green. In the crystal structure, the clashing region on the partner is disordered. In the MD simulations, though flexible, this region remains mostly ordered, causing additional deformation compared to the crystal structure. Illustrating the differences in strain localization among trajectories shown in panel (b), in the first two trajectories (red and yellow), the switch conformation aligns with the crystal structure and the partner deforms more; in the third trajectory (light blue), the partner conformation aligns with the crystal structure and the switch deforms instead. (Right) experimentally measured DEER distance distribution of the ternary complex (green with shaded 95% confidence interval) and distance distributions simulated from the crystal structure (gray line) or MD trajectories (dashed lines, colors correspond to the conformations shown at left). The distance distribution simulated from the crystal structure aligns with the left peak in the experimental distance distribution, whereas the distance distributions simulated from the MD trajectories span the experimental distance distribution, suggesting that these trajectories more fully sample the space of ternary complex dynamics.

**Extended Data Fig. 4:**
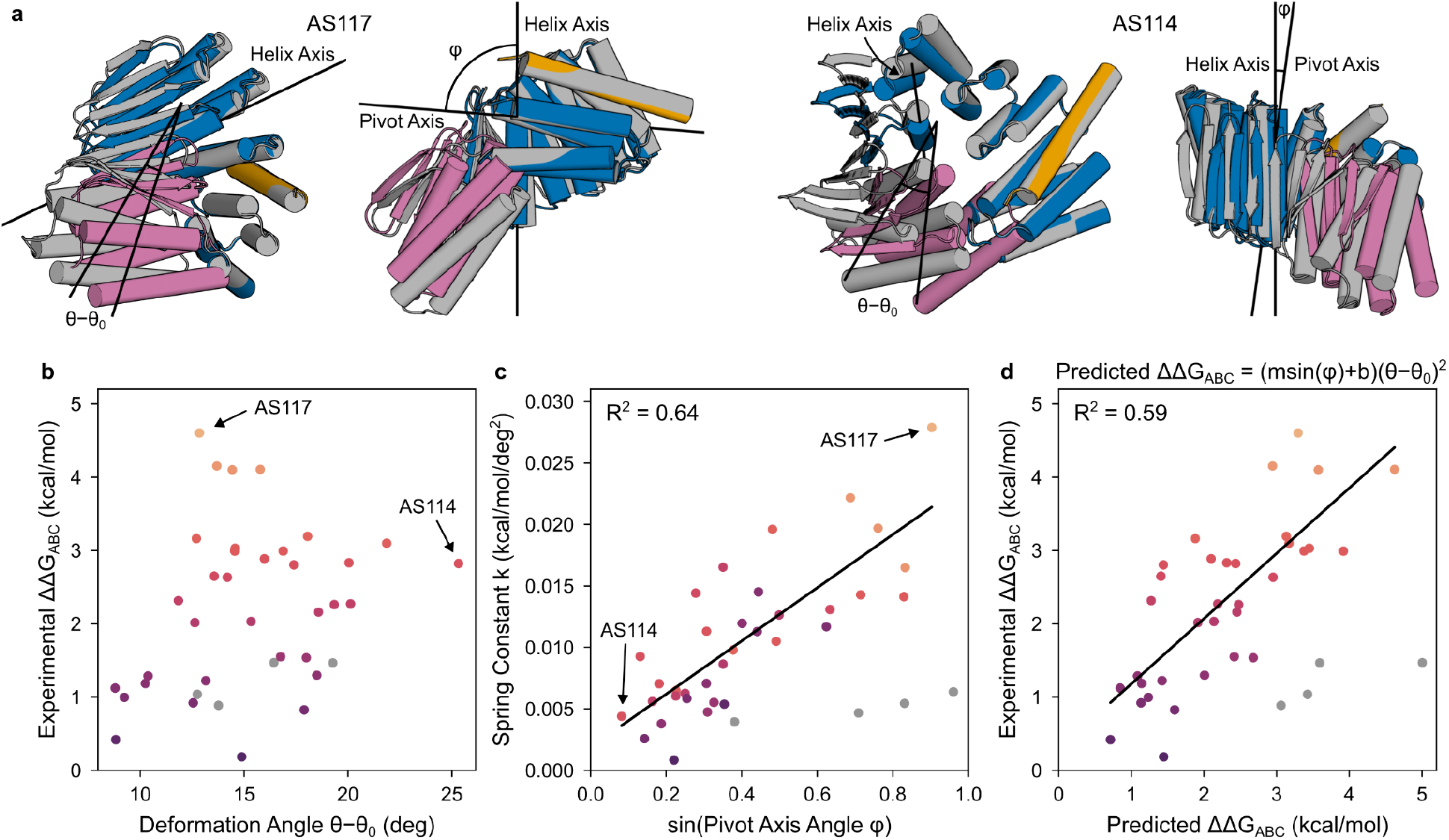
Modeling strain energy in the ternary complex. **a**, Differences in ternary complex geometry for a fast AS1 variant (AS117, left) and a slower variant which yet deforms more in the ternary complex (AS114, right). Design models of host-effector complex in state Y (blue and orange) aligned to the partner (pink) showing the allosteric clash, and (gray) AF2 predictions of the ternary complex aligned to the switch showing how the clash resolves through global strain. To model the strain energy from these structures using Hooke’s law, we measured the global deformation by the angle (*θ*−*θ*_0_) the partner pivots around some axis (the “pivot axis”) to move from its clashing position to its AF2-predicted strained position. The binder helix in the interface with the partner tends to be positioned near the centerpoint of the deformation (the point around which the centroid of the partner pivots), and the structure of this helix is the same for all variants, so we used the axis of this helix (the “helix axis”) to approximate the orientation of the deforming secondary structure elements. For each variant shown, the left view places the pivot of the partner within the plane of the page (so the pivot axis is normal to the page), and the right view places the helix axis and the pivot axis both within the plane of the page. In the left view, lines are drawn from the pivot axis through the centroids of the clashing and strained partners; the angle between these lines is the angle of global deformation (*θ*−*θ*_0_). The right view shows the angle between the helix axis and the pivot axis (*φ*); this is large for the fast variant and small for the slow but highly deforming variant. **b**, Lack of correlation between predicted magnitude of deformation and the experimental strain energy of the ternary complex for a set of host designs (AS1, AS2, AS5, AS7, and the AS1 variants, plotted as circles colored by their experimental strain energy except for AS101, AS115, AS119, and AS120 which were left out of the analysis in (c) and (d) and are colored gray—34 designs in total). The strain energy can be estimated from the observed accelerated off-rate as follows.

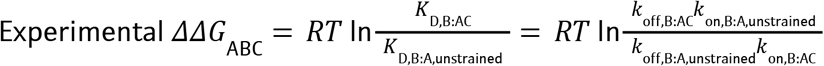

Making the approximation that *k*_on,B:AC_ = *k*_on,B:A,unstrained_ and does not vary with *ΔΔG*_ABC_ (Fig. S2) simplifies this expression.

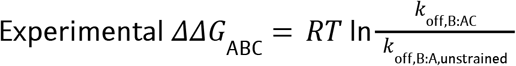

The accelerated off-rate *k*_off,B:AC_ is assumed to be the maximum off-rate observed in the facilitated dissociation experiments (Fig. S10). Since the main partner-binder interface does not change across variants, the off-rate of the unstrained interface *k*_off,B:A,unstrained_ should be a constant, *k*_base_, with one exception: in some variants, the switch fusion may form additional stabilizing interactions with the partner, reducing the base off-rate *k*_off,B:A_ (also measured in the facilitated dissociation experiments). These interactions can likely still form in the strained ternary complex to reduce the accelerated off-rate *k*_off,B:AC_ by the same factor. Thus, the smaller of *k*_base_ and *k*_off,B:A_ is used for *k*_off,B:A,unstrained_. For *k*_base_, a value of 2e−4 s^−1^ was used because it gave the best correlation between predicted and experimental *ΔΔG*_ABC_ described in panel (d). Notably, this value is quite close to the partner off-rate from the unhindered binder fusion LHD101B4, 9e−5 s^−1^ (Fig. S7). **c**, Linear correlation between the “spring constant” *k* (which relates the experimental strain energy to the magnitude of the predicted deformation) and the perpendicularity of this deformation to the secondary structure elements in the complex for this set of host designs (circles colored as in (b)). A linear regression on the colored points is plotted as a black line. The perpendicularity is computed as the sine of the angle between the pivot axis and the helix axis, and the spring constant is computed using Hooke’s Law as follows.

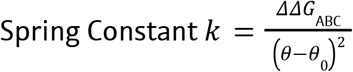

This relationship suggests that deforming in a stiff direction (against rather than around helices^33^) deforms the partner interface in a more destabilizing direction or better localizes strain to the partner interface instead of distributing the strain throughout the entire protein. **d**, Agreement between the experimental strain energy and the strain energy estimated entirely from the predicted structure of the strained ternary complex using Hooke’s Law for this set of host designs (circles colored as in (b)). A linear regression on the colored points is plotted as a black line. The spring constant was assumed to depend linearly on the deformation perpendicularity to the secondary structure elements, and the parameters of this correlation (m and b) were varied to fit the following expression to the experimental strain energies.

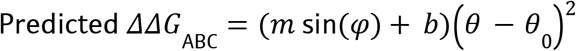

<u>Sources of Error in this Analysis</u> First, in this simple energetic model of facilitated dissociation described here and in Fig. S2, when the effector binds, the energy of the partner dissociation transition state will decrease by the binding energy of the effector. When the effector and partner are uncoupled, the energy of the ternary complex intermediate will decrease by the same amount so the activation barrier for partner dissociation will not change. When strain between the partner and effector is incorporated into the ternary intermediate, its energy will decrease less upon effector binding, reducing the activation barrier for partner dissociation. This simple energetic model thus assumes that the activation barrier for partner dissociation is directly related to the global energy of the ternary complex. In reality, acceleration of partner dissociation will arise from local deformation at the partner interface^89^. Thus, put differently, the simple energetic model assumes that the global deformation is smoothly distributed across the entire protein. This assumption allows us to directly relate the global deformation predicted by AF2 to the local deformation at the partner interface measured by the accelerated partner off-rate. In reality, the deformation is likely not smoothly distributed, and this may at least partially explain the imperfect correlation between the predicted and experimental strain energy of the ternary complex. This is exemplified by how the partner dissociates more slowly from the AS1 ternary complex with the peptide effector than with the 3hb effector (Fig. 2e,f). Despite forming the same interactions and causing the same conformational change as the 3hb, the more deformable peptide appears to less effectively localize strain to the partner once in the ternary complex. Uneven distribution of strain in the ternary complex is also one possible mechanism underlying unidirectional competition in facilitated dissociation systems. Second, our model estimating the protein stiffness anisotropy using the orientation of the secondary structure elements is likely oversimplified. Proteins have finer levels of structure than the orientation of their secondary structure elements. Much like a normal mode analysis, a more sophisticated model could estimate local spring constants for smaller regions of the protein, then compute a strain energy for each region from its predicted deformation. Third, inaccuracies in the AF2 predictions of the strained ternary complex could cause much of the variation between the experimental and predicted strain energies. Clear cases of this, variants AS101, AS115, AS119, and AS120 (gray) which do not follow this correlation were left out of this analysis. We hypothesize that these variants can adopt an alternate, less-strained ternary complex conformation than the high-energy conformation predicted by AF2.

**Extended Data Fig. 5:**
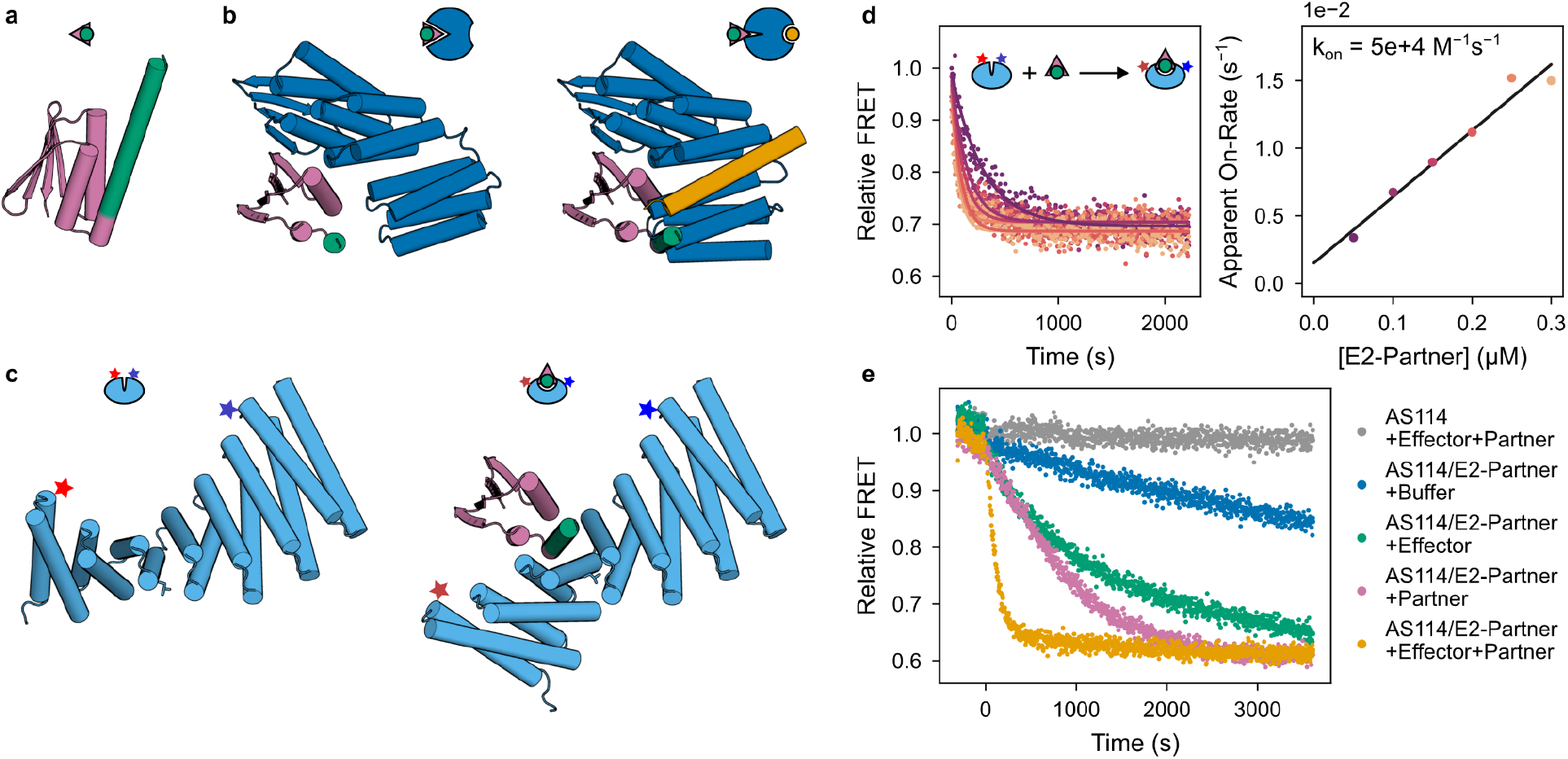
Construction and characterization of the chain reaction. **a**, Design model of E2-partner, comprising the partner LHD101A (with mutations R43V and V69Q) fused to the effector peptide “E2” (cs201B) for hinge cs201. E2 is colored green and LHD101A is colored pink. **b**, Design models of E2-partner (green/pink) bound to AS114 (blue) in state X showing no clash (left) and in state Y with the effector peptide (orange) showing a strong clash (right). **c**, Design models of the reporter hinge “H2” (cs201F with mutation E249L (sticks) which increases E2 on-rate and labeled with Alexa Fluors 555 and 647 at positions indicated by stars) in state X (left) and in state Y with E2-partner (right). AS114 and H2 would massively overlap if simultaneously bound to E2-partner, so their binding should be mutually exclusive: AS114 should cage E2 until its release by the effector. **d**, E2-partner and H2 association rate, measured by a change in FRET efficiency due to the conformational change in H2 upon binding. (Left) FRET time courses (normalized to the initial signal) with varying concentrations of E2-partner and 5 nM H2; data (circles) fit with single exponentials (lines). (Right) apparent on-rates plotted against E2-partner concentration (circles) and a linear fit. The value of the association rate constant (5e+4 M^−1^s^−1^) is higher than the reported value (4.5e+3 M^−1^s^−1^) for the original hinge cs201F with effector cs201B, suggesting that mutation E249L on H2 biases its conformational pre-equilibrium toward state Y to increase the apparent association rate. **e**, Additional data for the kinetically governed chain reaction shown in Fig. 4b. In the gray control time course, 500 nM AS114 was added to 20 nM H2, then 1 μM effector and 6 μM partner was added at time 0, showing that none of these components bind to H2 to cause a change in FRET signal. In the other time courses, preincubated 500 nM AS114 and 250 nM E2-partner was added to 20 nM H2, then buffer (blue), 1 μM effector (green), 6 μM partner (pink), or both (orange) were added at time 0. A baseline drift (obtained from 500 nM AS114 after adding 20 nM H2 then at time 0 adding buffer) was subtracted from each time course, and time courses were normalized to the initial signal. The chain reaction proceeds faster when just excess partner is added, likely due to blocking rebinding of E2-partner to AS114 after transient dissociation, but this effect is insufficient to achieve full acceleration. The chain reaction also proceeds faster when just effector is added, but likely due to transient rebinding of E2-partner to re-form the strained ternary complex, this also does not achieve full acceleration. Adding both effector (to accelerate E2-partner dissociation from AS114) and excess partner (to prevent E2-partner rebinding to AS114) is required to fully accelerate the chain reaction. Note that if a single-chain effector is desired to fully accelerate the chain reaction, the effector and partner could be flexibly fused into a single construct. Such a multivalent effector would be reminiscent of CITED2, whose multivalency enables rapid and unidirectional competition against HIF-1α^90^.

**Extended Data Fig. 6:**
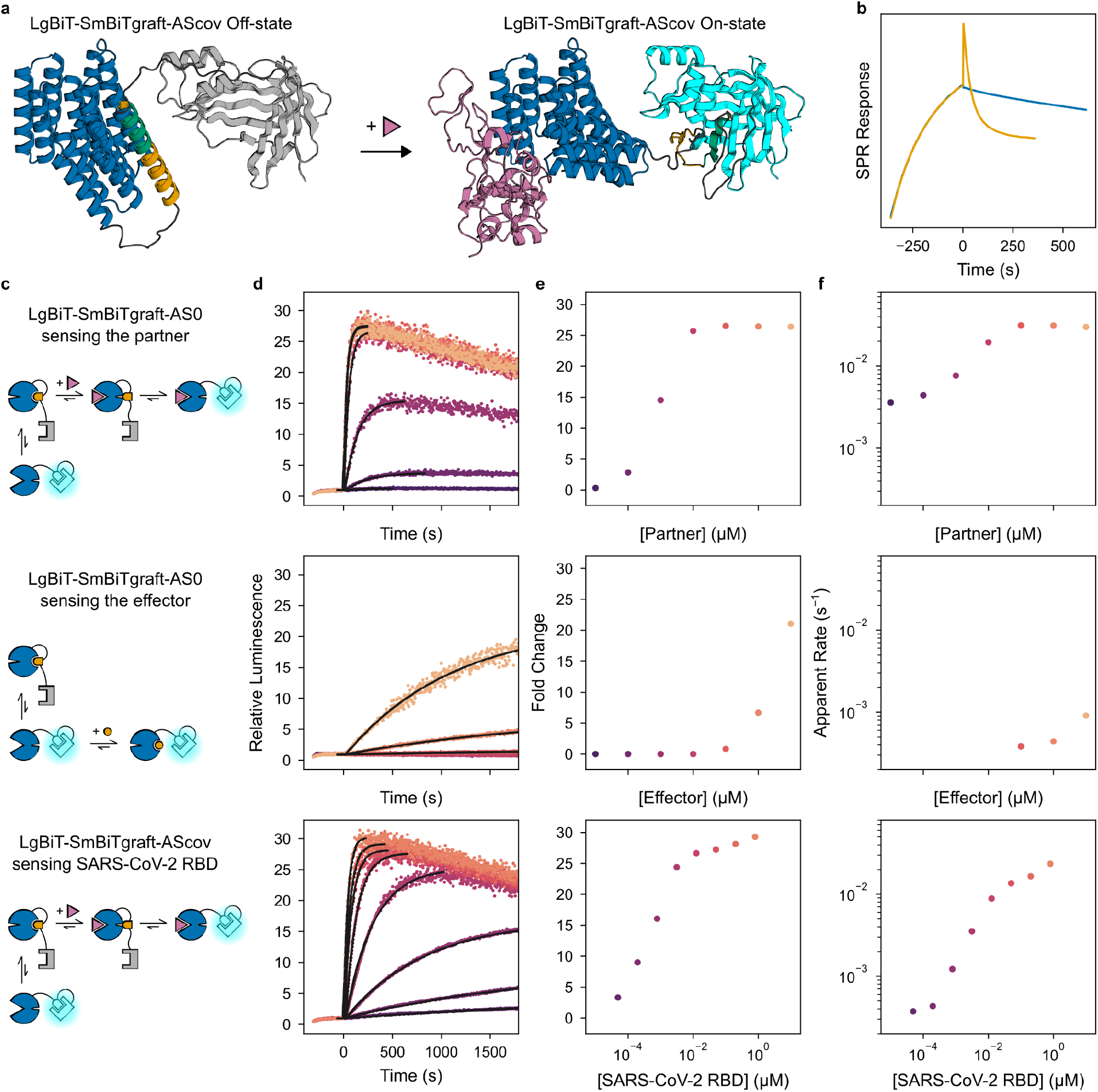
Construction and characterization of rapid sensors. **a**, Structural model of the best SARS-CoV-2 sensor construct, comprising AScov (blue), the SmBiTgraft peptide with the effector (orange) and grafted SmBiT (green), LgBiT (gray or cyan), and flexible linkers (black). **b**, SPR data showing sfGFP-SmBiTgraft binding to AS0 (blue and orange, association phase), slow subsequent dissociation in the absence of partner (blue), and rapid subsequent dissociation upon addition of 10 μM partner (orange) caused by rapid partner binding to form a transient ternary complex, causing the spike at the beginning of the dissociation phase. **c–f**, Rapidly sensing the partner through a facilitated dissociation mechanism (top), slowly sensing the effector limited by the slow base exchange rate of SmBiTgraft between binding AS0 and LgBiT (middle), and rapidly sensing the SARS-CoV-2 RBD with facilitated dissociation (bottom). **c**, Schematics showing the mechanism of sensing. **d**, Luminescence time courses (normalized to the initial signal) of 10 pM sensor construct then at time 0 adding varying concentrations of analyte; data (colors) fit (black) with single exponentials up to the maximum signal for time courses which showed appreciable signal increase. In some time courses, signal slowly decreases due to depletion of luciferase substrate. **e**, Luminescence signal fold change plotted against analyte concentration. **f**, Sensor response rate plotted against analyte concentration for time courses which showed appreciable signal increase.

**Extended Data Fig. 7:**
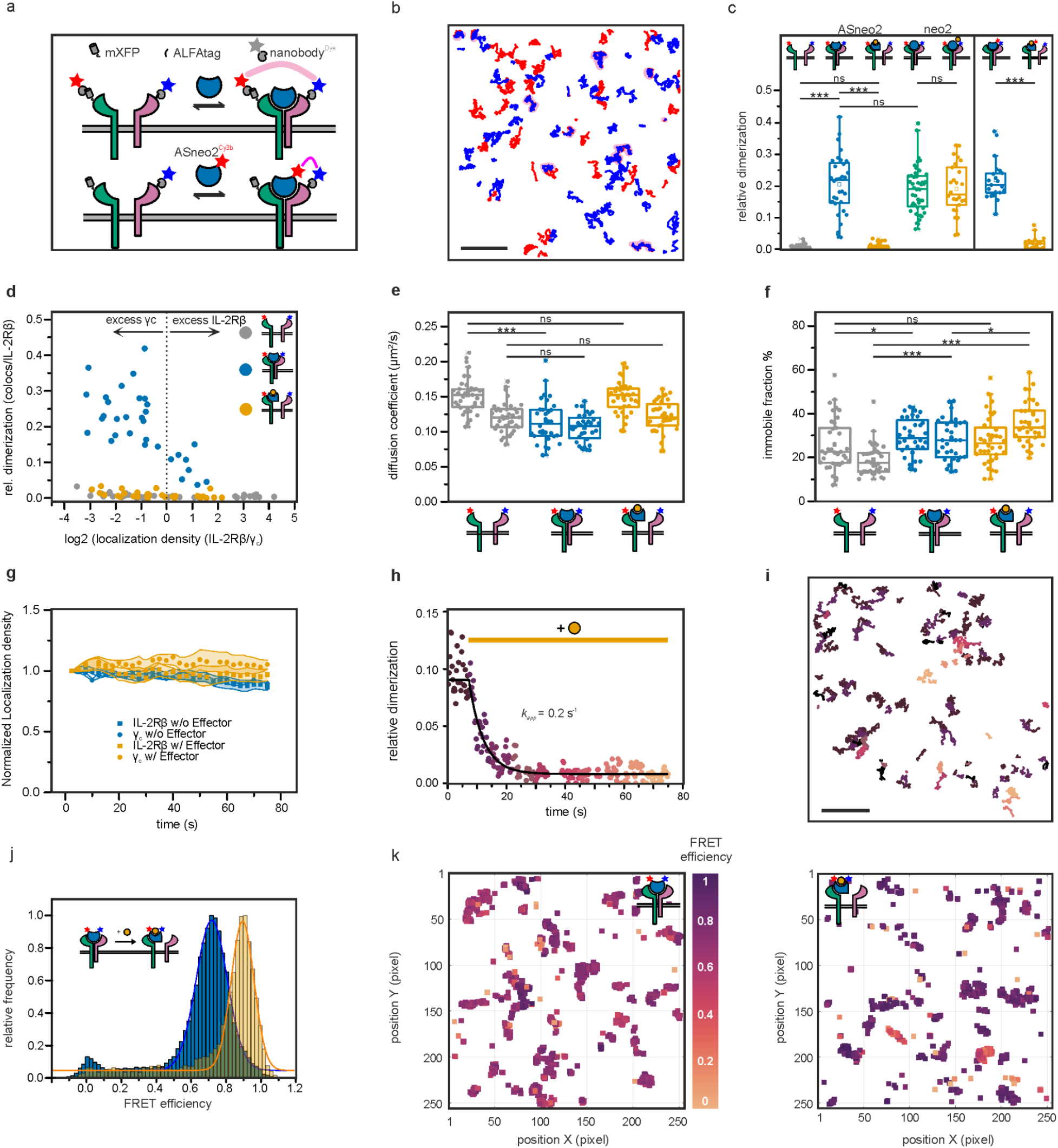
Detailed functional characterization of ASNeo2. **a**, Schematic depiction of labeling strategy for single molecule tracking experiments **b**, Single molecule trajectories of IL-2Rβ (red), γ_c_ (blue) and ASNeo2-induced heterodimers (magenta). **c**, Data from Fig. 5e as boxplots to display datapoint variation including Neo2 (+/-effector) (green and orange, left side) and single molecule tracking experiments with labeled ASNeo2 and γ_c_ (right side). Sample sizes and independent repeats are: unstimulated: 37 and 3; ASneo2: 32 and 3; ASneo2 + Effector: 33 and 3; neo2: 44 and 4; neo2 + Effector: 37 and 3; labeled ASneo2: 21 and 2; labeled ASneo2 + Effector: 18 and 2. **d**, Relative dimerization in relation to receptor cell surface density ratio indicates that high dimerization data variance is caused by differing IL-2Rβ to γ_c_ ratios at the plasma membrane. Even at high γ_c_ excess, effector-bound ASNeo2 shows no residual affinity for γ_c_. **e**, Diffusion properties of IL-2Rβ and γ_c_ are reverted to the ground state after addition of effector. **f**, Immobile particles are increased upon stimulation, but not decreased after effector addition, potentially indicating receptors internalizing in membrane proximal endosomes. For e and f, the left box always corresponds to IL-2Rβ and the right one to γ_c_. Sample sizes for d–f are as in **c. g**, Normalized localization density over time confirms minimal single molecule bleaching in long term single molecule tracking experiments. Sample sizes and independent repeats are: without Effector: 5 and 5; with Effector: 3 and 3. **h, i**, Dissociation of ASNeo2-induced IL-2Rβ/γ_c_ dimers at the cell surface upon addition of 10 μM effector as detected by time-lapse single-molecule co-tracking (h) with color-coded corresponding co-trajectories (i). **j, k**, Conformational change of ASNeo2 bound to the cell surface receptor probed by smFRET. **j**, FRET efficiency histograms for ASNeo2 E4C/K211C labeled with Cy3B and ATTO643 bound to cells expressing IL-2Rβ and γ_c_ in the absence (blue) and presence (yellow) of 10 μM effector. Sample sizes are: without Effector: 7; with Effector: 5. **h**, smFRET co-localizations of one individual cell before the effector was added (left) and after it was added (right) color-coded for FRET efficiency, highlighting the observation of individual molecules. Statistics for c, e and f were performed using two-sided two-sample Kolmogorov–Smirnov tests (not significant (NS), * p < 0.05, *** p < 0.001). Boxplots show the distribution of the dataset, highlighting the median, quartiles, and outliers, with whiskers extending to the range limits. Scale bar in b and i: 5 μm.

**Extended Data Fig. 8:**
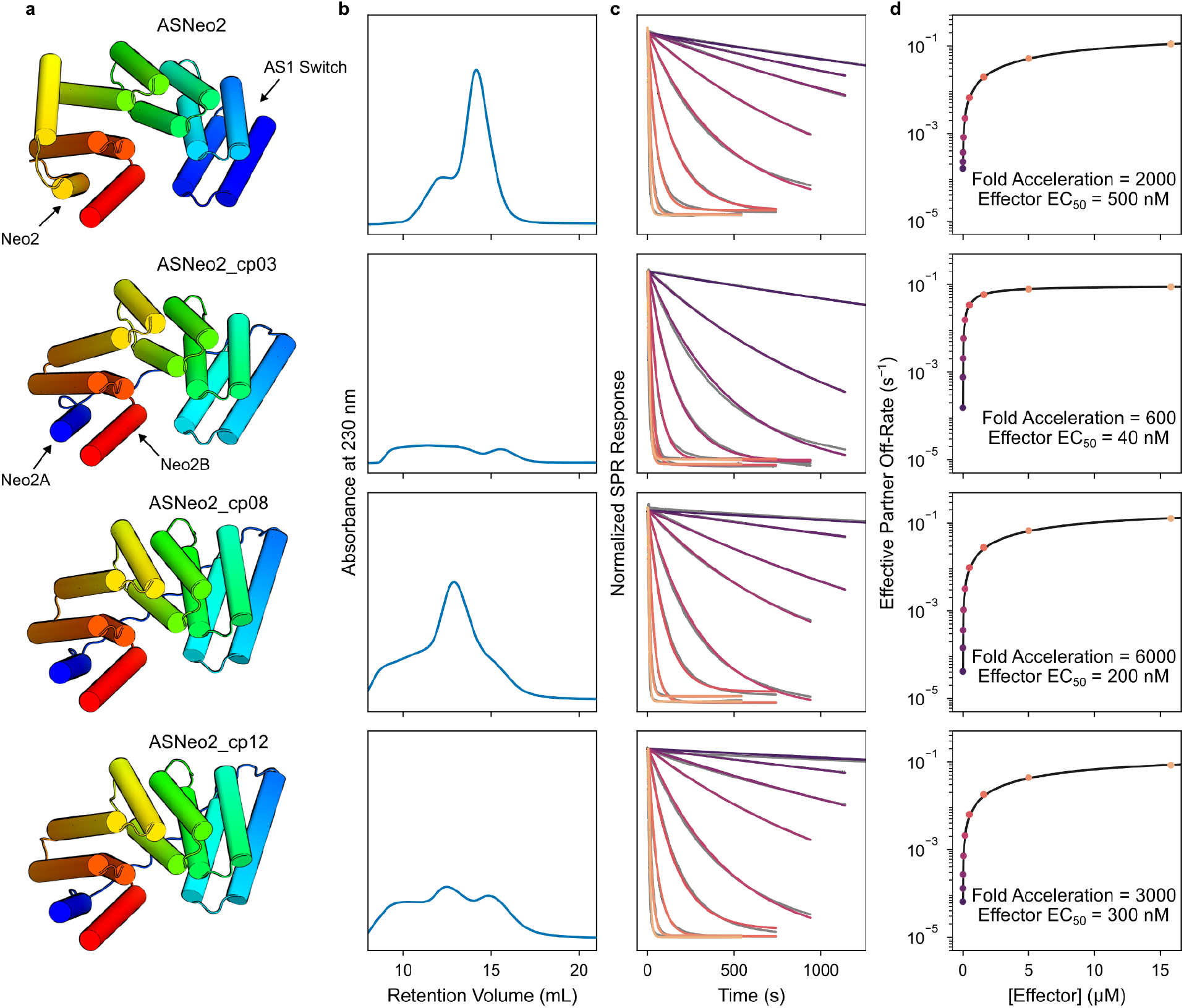
Characterization of cyclic permutations of ASNeo2. **a**, Design models of ASNeo2 and selected cyclic permutations in state X, rainbow-colored from N-terminus (blue) to C-terminus (red) to illustrate the protein topology. In ASNeo2, the switch is at the N-terminus and Neo2 is at the C-terminus. In the cyclic permutations, though the relative positions of the switch and Neo2 changes minimally, the switch is in the middle of the protein, part of Neo2 is at the N-terminus, and the other part is at the C-terminus. This way, the regulatory switch cannot degrade without also breaking Neo2. **b**, SEC purifications performed on a Superdex 200 Increase 10/300 GL column. The cyclic permutations are prone to aggregation during expression, but distinct monomer peaks can be picked out. **c**, Fast effector concentration–dependent dissociation of γ_c_ from the ASNeo2-IL-2Rβγ_c_ complex upon addition of peptide effector. Data (gray) fit (colors) as described in methods (neglecting the accumulation modeling because accumulation on the SPR surface was negligible with these proteins). **d**, Effective partner off-rates computed from the model fit by ln(2)÷{half-time of γ_c_-host interaction} plotted against effector concentration (circles) and fit with hyperbolic equations (black lines).

