## Supplementary Information for "Design of facilitated dissociation enables control over cytokine signaling duration"

**Nomenclature**

“Host” refers to the switch-binder fusion with allosterically coupled partner and effector binding sites.

“Partner” refers to the protein which binds tightly to the host but can be rapidly kicked off.

“Effector” refers to the protein/peptide which binds to the host at a different site to cause a conformational change in the host which destabilizes partner binding.

“Binder” refers to the component of the host designed to bind to the partner.

“Switch” refers specifically to the effector-responsive conformational switch component of the host proteins, and we avoid using “switch” to refer to the entire host protein. This way, we distinguish between designing switches (multi-state design) and designing hosts (fusing switches to binders).

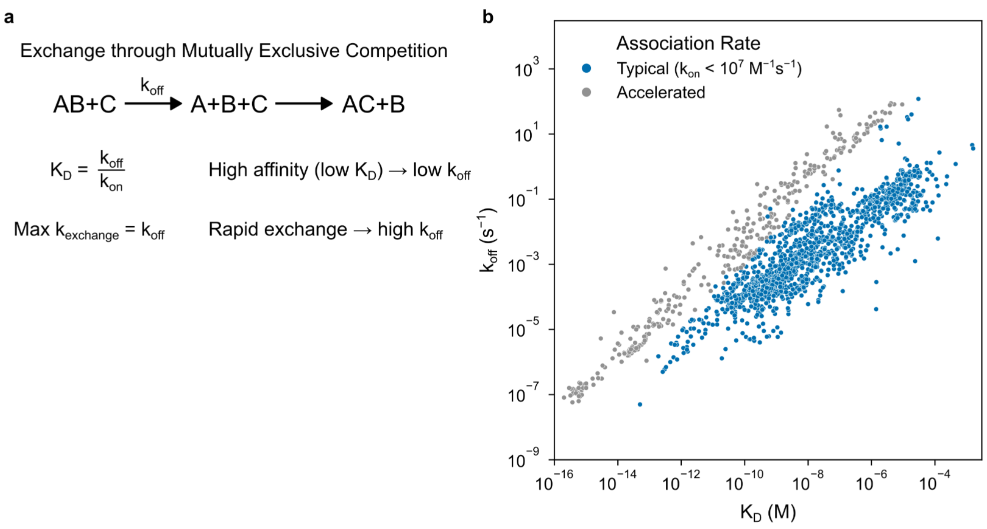


**Fig. S1: Relationship between affinity and exchange rate.** **a**, For protein-protein interactions with typical diffusion-limited association rates in the $10^5$–$10^6$ $M^{-1}s^{-1}$ range, high affinity ($K_D < 1$ nM) requires low off-rates ($k_{off} < 10^{-3}$ $s^{-1}$). This precludes rapid exchange through mutually exclusive competition, which requires high off-rates. **b**, Off-rate vs affinity for a set of natural and mutant proteins with typical on-rates (blue) and high on-rates (gray) depicting this tradeoff. Data obtained from the SKEMPI database[1].

High affinity and rapid exchange could be simultaneously achieved in binary interactions with high on-rates ($k_{on} > 10^7$ $M^{-1}s^{-1}$)[2], but these are usually caused by long-range electrostatic attraction and are not a general feature of protein-protein interactions. Although electrostatically accelerated association has been engineered into proteins[3], this is not possible in general with native targets because it requires patches of complementary surface charge on both binding partners.

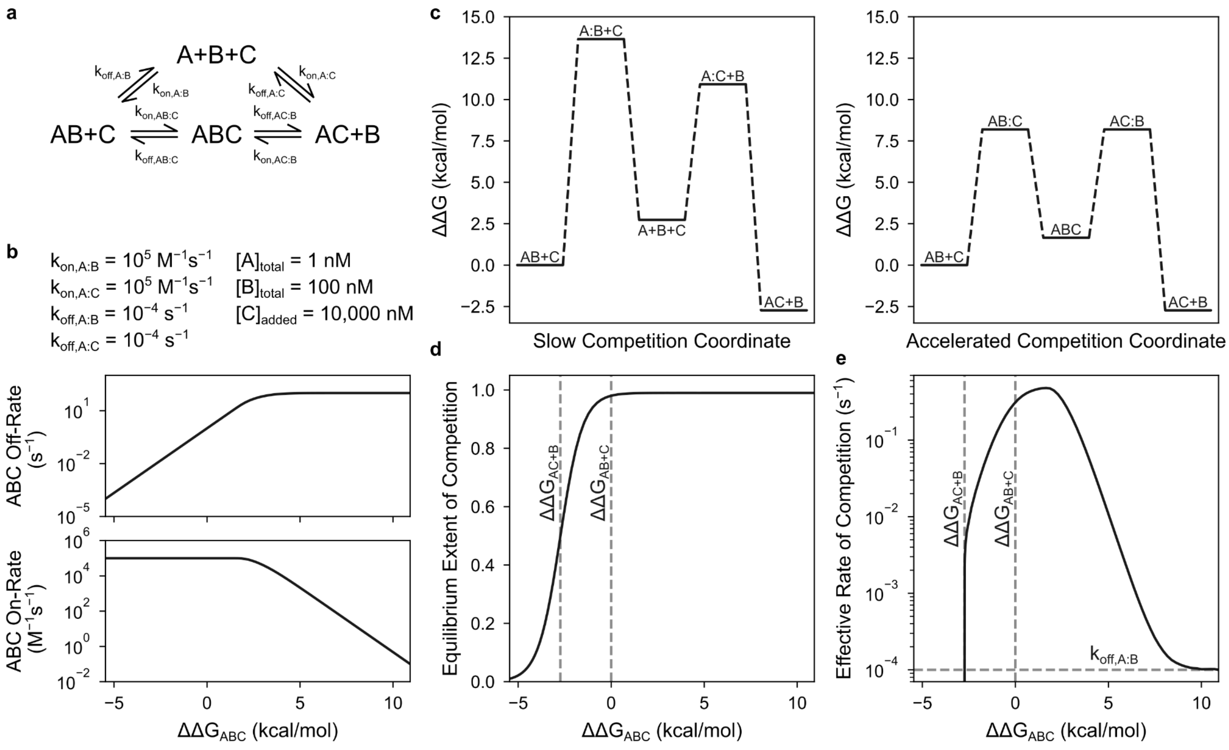


**Fig. S2: Dependence of facilitated dissociation on the energy of the ternary intermediate. a**, Kinetic model of a facilitated dissociation system. We modeled the dynamics of a facilitated dissociation process for a range of ternary intermediate energies, simulated in this model by pre-equilibrating A and B then adding excess C. **b**, Model parameters used to simulate facilitated dissociation. We fixed the binary interaction rate constants, and we varied the ternary interaction rate constants to set the ternary intermediate energy. The plots show the relationships we chose between the ternary intermediate energy and the ternary interaction rate constants. These hold with physical intuition: low to moderate increases in ternary intermediate energy primarily affect dissociation rates, whereas high increases in ternary intermediate energy additionally affect association rates. **c**, Energy diagrams of slow mutually exclusive competition (left) and facilitated dissociation (right). Colons denote binding transition states between components. Binding C reduces the energy of the entire system, including the B dissociation transition state. Frustration between B and C causes the ternary intermediate energy to reduce less, lowering the barrier for B dissociation. **d**, Dependence of the equilibrium extent of the competition reaction on the ternary intermediate energy. If the ternary intermediate energy is too low (too little strain), the ternary complex becomes the dominant equilibrium state and B dissociation is not favored. **e**, Dependence of the effective rate of competition on the ternary intermediate energy. The effective rate is defined as ln(2)÷{time to 50% extent of reaction} (this is undefined when $\Delta\Delta G_{ABC} < \Delta\Delta G_{AC+B}$ because the final extent of reaction is less than 50%). The rate of competition peaks at an optimal ternary intermediate energy, then recedes to the basal rate of mutually exclusive competition as the ternary intermediate becomes increasingly energetically inaccessible.

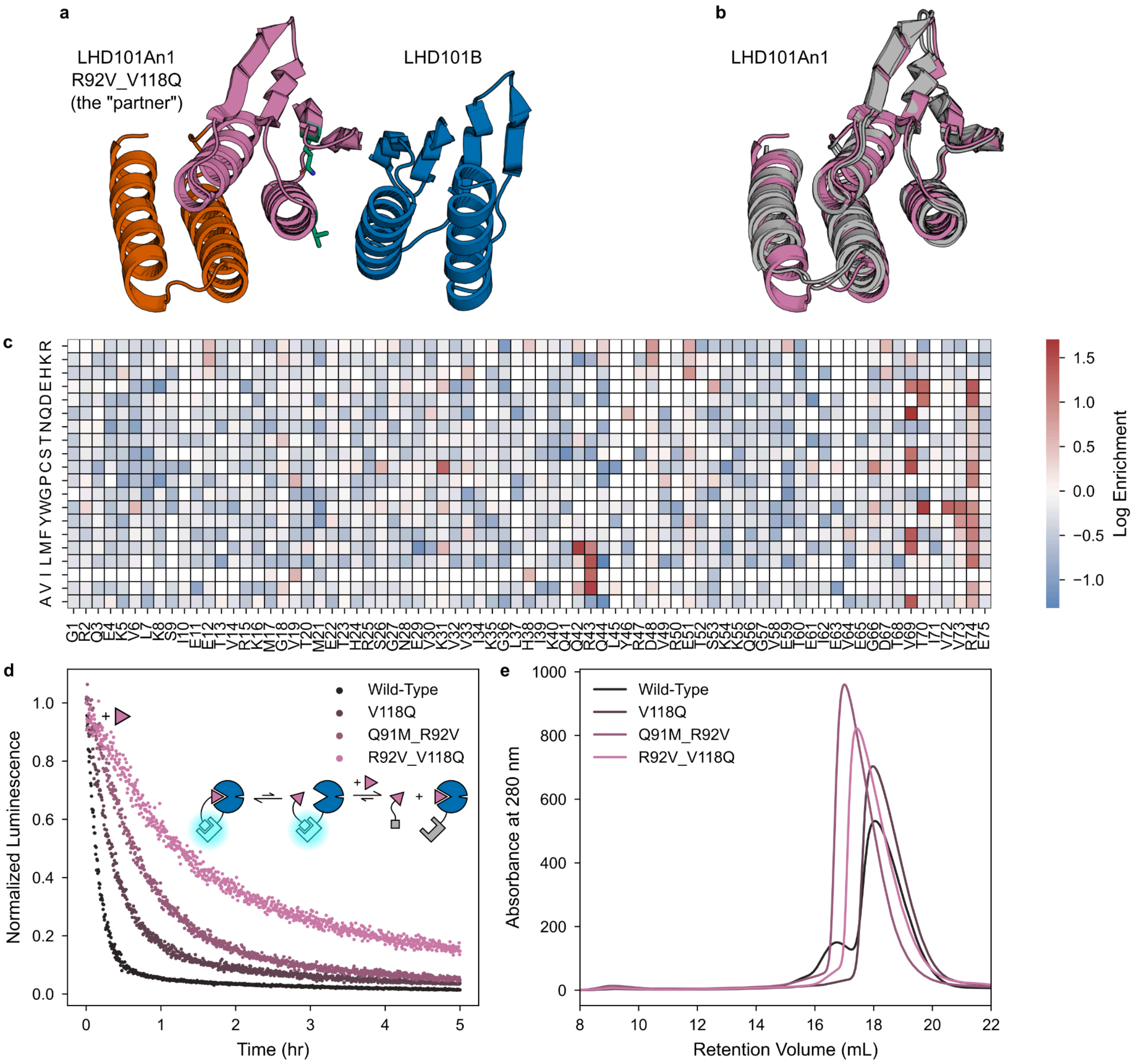


**Fig. S3: Design and characterization of the partner. a**, Design models of the partner (pink/red) and binder (blue) in complex. Starting from LHD101A (pink), we built additional structure (red) to increase opportunities for steric clashing, resulting in the partner protein "LHD101An1." Unless otherwise stated, "partner" refers to LHD101An1 with affinity-enhancing mutations R92V and V118Q (green). **b**, Crystal structures of LHD101An1 (gray) overlaid with the design model (pink). **c**, Site-saturation mutagenesis heatmap showing the enrichment over wild-type of each LHD101A point mutation displayed on yeast sorted against 10 nM LHD101B. Mutations Q42M, R43V, and V69Q on LHD101A correspond to Q91M, R92V, and V118Q on LHD101An1. **d**, Dissociation time courses of preincubated 20 nM LHD101An1-SmBiT mutants and 100 pM AS0-LgBiT after adding 20 µM LHD101An1, showing the double mutant R92V V118Q in particular greatly reduces the intrinsic partner-host off-rate. **e**, SEC purification runs of LHD101An1 mutants, performed on a Superdex 200 Increase 10/300 GL column with injection concentrations greater than 500 µM, showing these mutants remain well-behaved in solution.

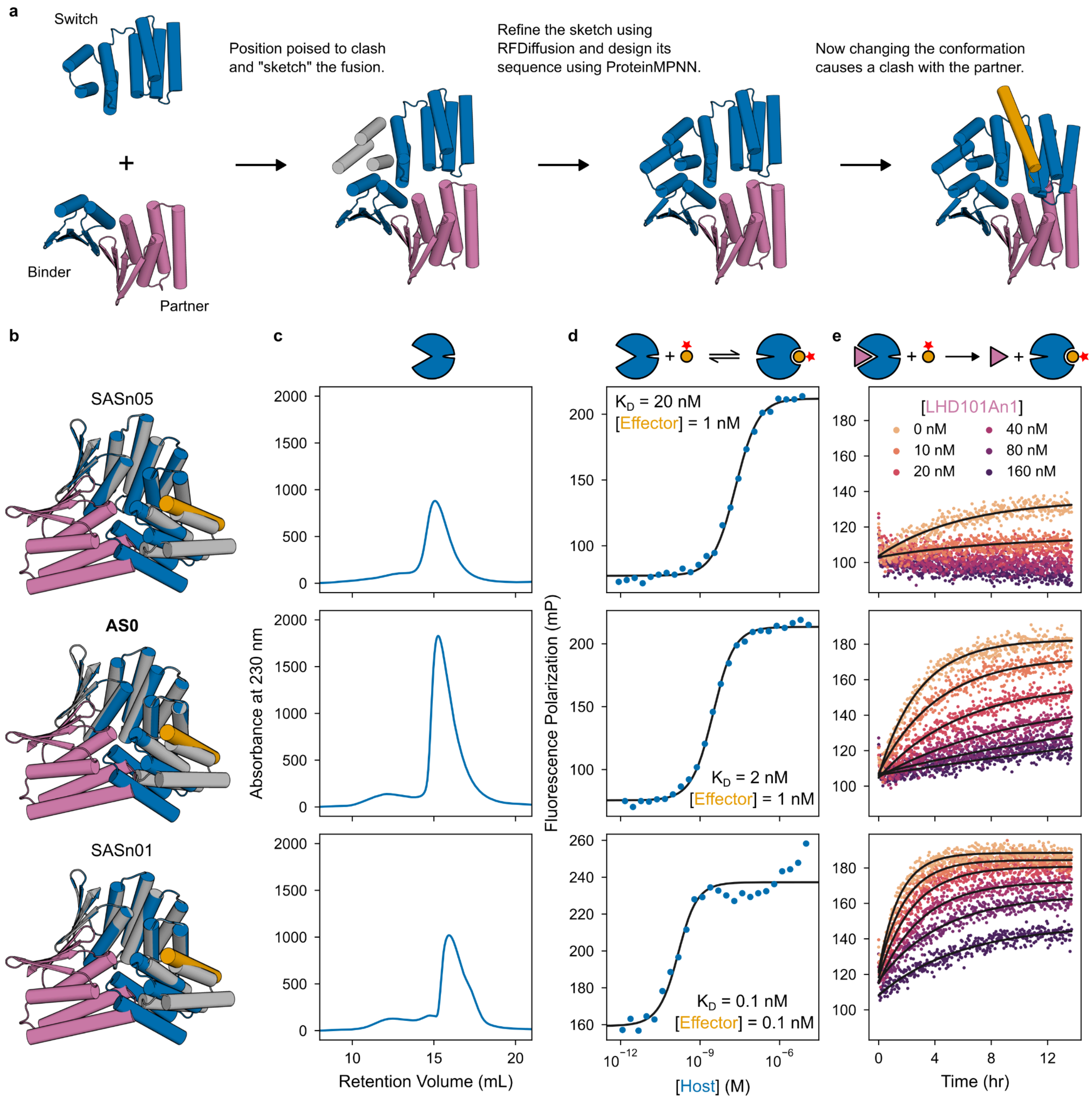


**Fig. S4: Negative allosteric coupling between partner and effector. a**, Approach to designing host proteins by fusing a partner binder and an effector-responsive switch. **b**, Design models of selected cs221-based host-effector complexes in state Y (blue and orange) aligned to the partner (pink) and to the host in state X (gray). **c**, SEC host purifications performed on a Superdex 200 Increase 10/300 GL column. **d**, Fluorescence polarization (FP) titrations with a constant concentration of TAMRA-labeled effector and varying host concentrations. Data (blue) fit with a standard binding isotherm (black). **e**, Association of 20 nM TAMRA-labeled effector to 20 nM host against varying concentrations of LHD101An1, showing competition between partner and effector binding. FP data (colored) fit with single exponentials (black).

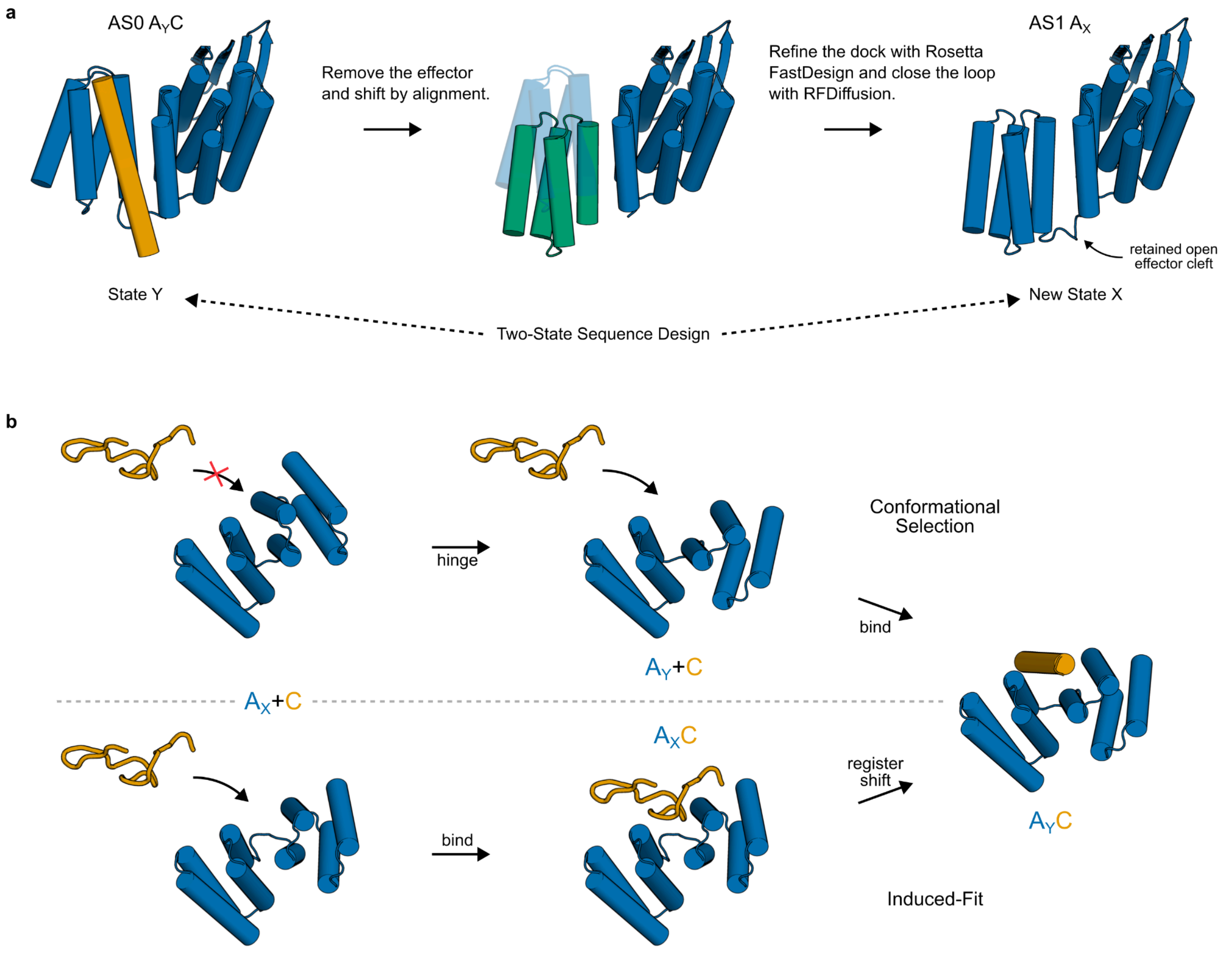


**Fig. S5: Design of induced-fit effector binding. a**, Approach to designing register-shift host proteins which retain an open cleft in state X. **b**, Comparison of effector binding mechanisms for the original hinges (top) and new register-shift switches (bottom). The closed state X of the original hinge blocks effector binding, so the hinge must change conformation before the effector can bind: a conformational selection mechanism. The open state X of the new switches could allow weak effector binding which triggers the conformational change: an induced-fit mechanism.

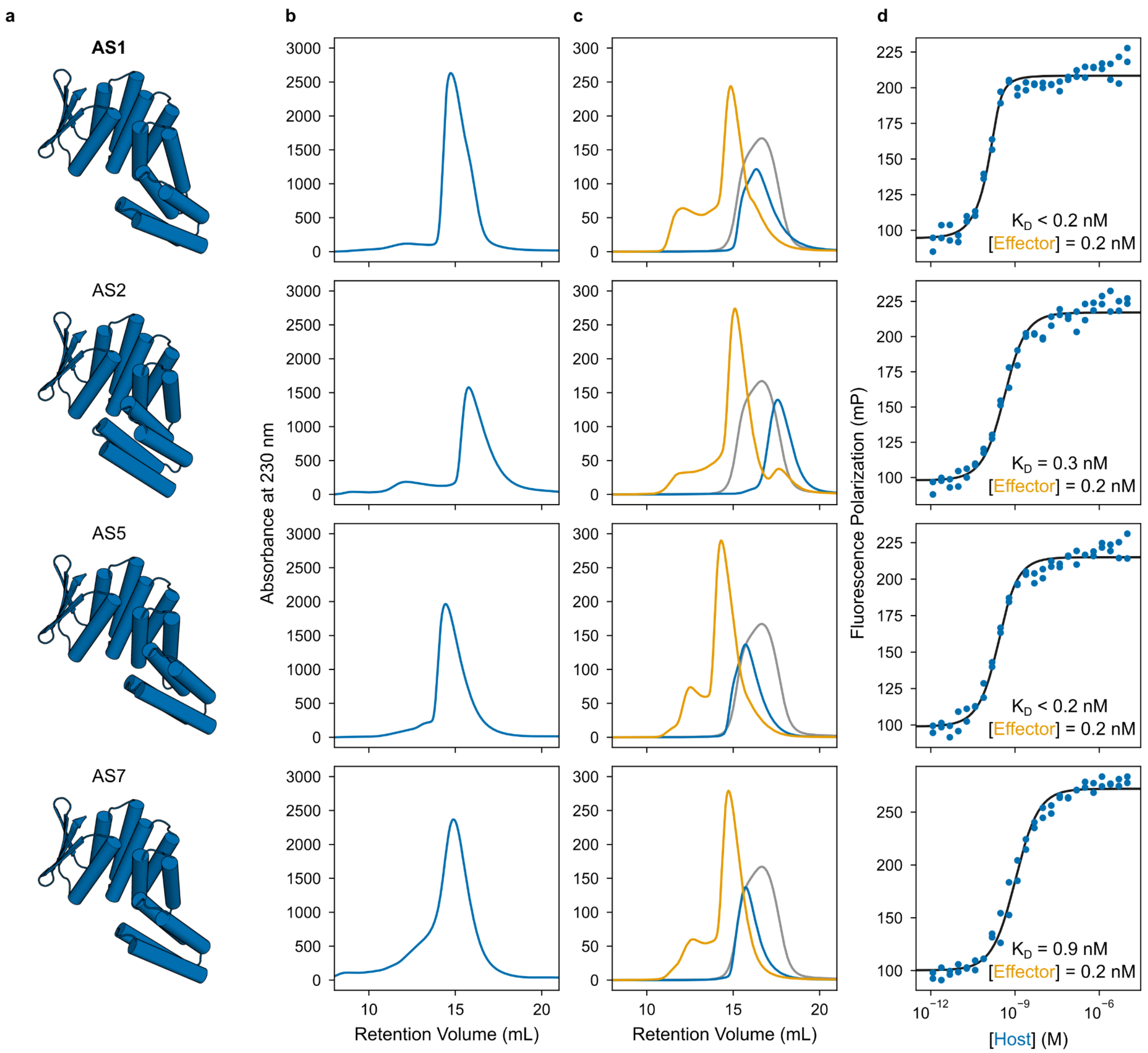


**Fig. S6: Initial characterization of register-shift host designs. a**, Design models of selected register-shift host proteins in state X, showing the diversity of the new state X. **b**, SEC host purifications performed on a Superdex 200 Increase 10/300 GL column. **c**, SEC binding experiments performed on a Superdex 200 Increase 10/300 GL column. All components were injected at 20 μM. The mixtures of host and sfGFP-tagged effector (orange) run larger than the hosts alone (blue) or sfGFP-effector alone (gray), indicating host-effector binding. **d**, FP titrations with a constant concentration of TAMRA-labeled effector and varying host concentrations. Data (blue) fit with a standard binding isotherm (black).

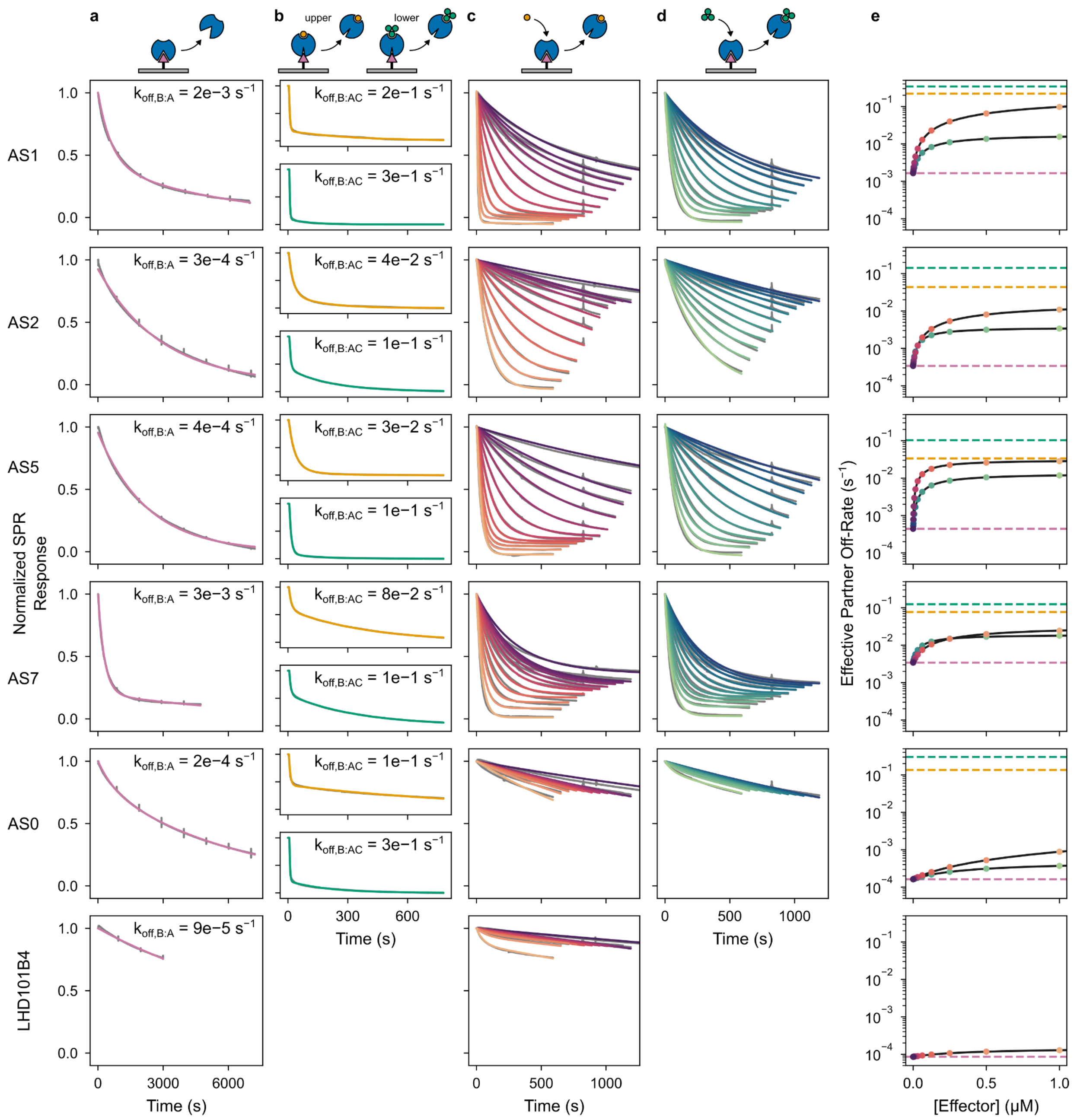


**Fig. S7: SPR characterization of facilitated dissociation in host designs. a**, Slow dissociation of the partner from the host in the absence of effector. Data (gray) fit with exponential decay functions (pink). For AS2 and AS5, single exponentials were used, whereas for the other designs, double exponentials were used to account for populations of host protein with different dissociation kinetics. For the double exponential fits, the reported dissociation rate constant, $k_{off,B:A}$, is the rate constant from the higher amplitude exponential in the fit. **b**, Faster dissociation of the partner from the ternary complex with effector. In each row, the top plot corresponds to the peptide effector and the bottom plot to the 3hb effector. Data (gray) fit with double exponential decay functions (orange for peptide, green for 3hb) to account for a population of partner-host complex lacking the effector. The reported dissociation rate constant, $k_{off,B:AC}$, is the higher of the two rate constants in the fit. **c** and **d**, Effector concentration–dependent dissociation of the partner upon addition of peptide (c) or 3hb (d) effector. Data (gray) fit (colors) as described in methods. **e**, Effective partner off-rates computed from the model fit by ln(2)÷{half-time of partner-host interaction} plotted against effector concentration (circles) and fit with hyperbolic equations (black lines). The orange circles correspond to the data with peptide effector from (c) and the green circles to the data with 3hb effector from (d). The pink line plots $k_{off,B:A}$ measured in (a), the orange line plots $k_{off,B:AC}$ with peptide effector measured in (b), and the green line plots $k_{off,B:AC}$ with 3hb effector measured in (b). With the peptide effector, the effective rate of the full facilitated dissociation pathway approaches the rate of partner dissociation from the ternary intermediate, whereas with the 3hb effector, the effective rate of the full facilitated dissociation pathway approaches a lower value. As observed for AS1 (Fig. 2), this likely corresponds to peptide binding through induced-fit and 3hb binding rate-limited by a slower conformational selection. At the top of **a**–**d**, cartoons show the arrangement of proteins relative to the SPR chip (gray). Only the experiments of panels (a) and (c) were performed for the LHD101B4 control. Also note that the data in gray is often hidden behind the colored fit curves.

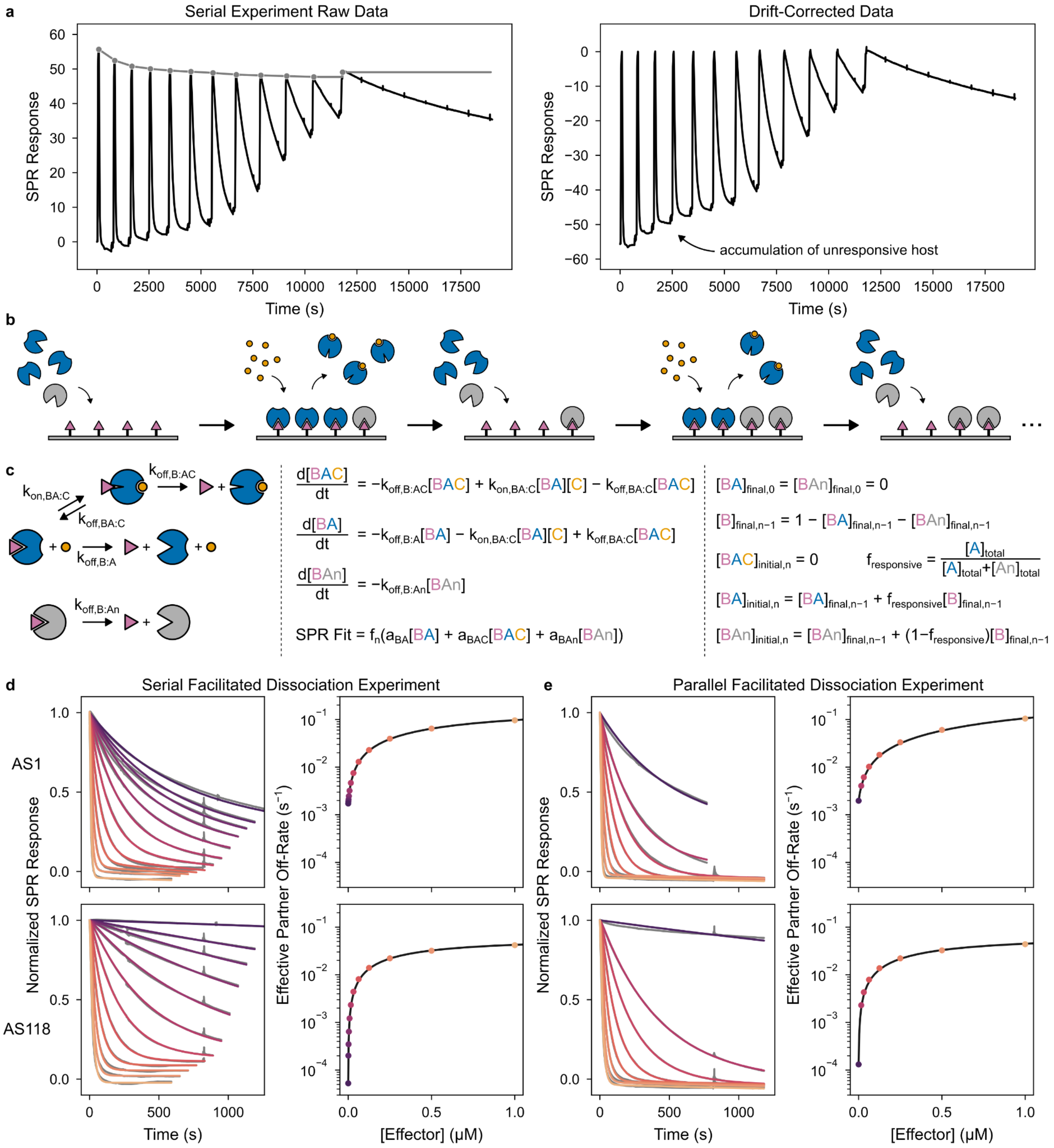


**Fig. S8: Fitting SPR facilitated dissociation data. a**, (Left) raw SPR data (black) of a serial facilitated dissociation experiment involving multiple cycles of associating host then dissociating it with varying concentrations of effector. (Gray) approximate baseline drift estimated by interpolating between the peaks of host association (the SPR response should be the same when the surface is saturated with host). (Right) SPR data corrected for baseline drift by subtracting the gray baseline approximation from the raw SPR data. After each cycle, the dissociation trace plateaus at increasingly higher SPR response values, likely corresponding to accumulation of a small population of host that is unresponsive to the effector. **b**, Cartoons showing how a small population of unresponsive host (gray, “An”) will accumulate on the surface after multiple facilitated dissociation cycles. **c**, Kinetic model fit to the facilitated dissociation experiment (see methods). (Left) cartoons depicting the state transitions which would affect the SPR response. (Middle) The system of differential equations corresponding to the kinetic model on the left, which can be fitted to the dissociation curve of each cycle. The bottom equation relates the concentrations of each state on the SPR surface to an SPR response. (Right) initial values for fitting this model to the dissociation curve of cycle *n*. **d** and **e**, (Left) fit data from serial (d) or parallel (e) facilitated dissociation experiments on AS1 (top row) or AS118 (bottom row). Data shown in gray and fits in colors. In the parallel facilitated dissociation experiment, each effector concentration is tested on a fresh SPR surface, minimizing accumulation of unresponsive host. Not needing to account for this, the model used to fit parallel facilitated dissociation data can be simpler. (Right) Effective partner off-rates computed from the model fit by ln(2)÷{half-time of partner-host interaction} plotted against effector concentration (circles, colors correspond to the fit dissociation traces in the left plots) and fit with a hyperbolic equation (black line). The partner dissociation kinetics obtained from fitting the serial and parallel facilitated dissociation experiments agree within 4-fold, indicating that the serial experiment provides fairly accurate measurements of the partner dissociation kinetics despite requiring a more complex model to fit.

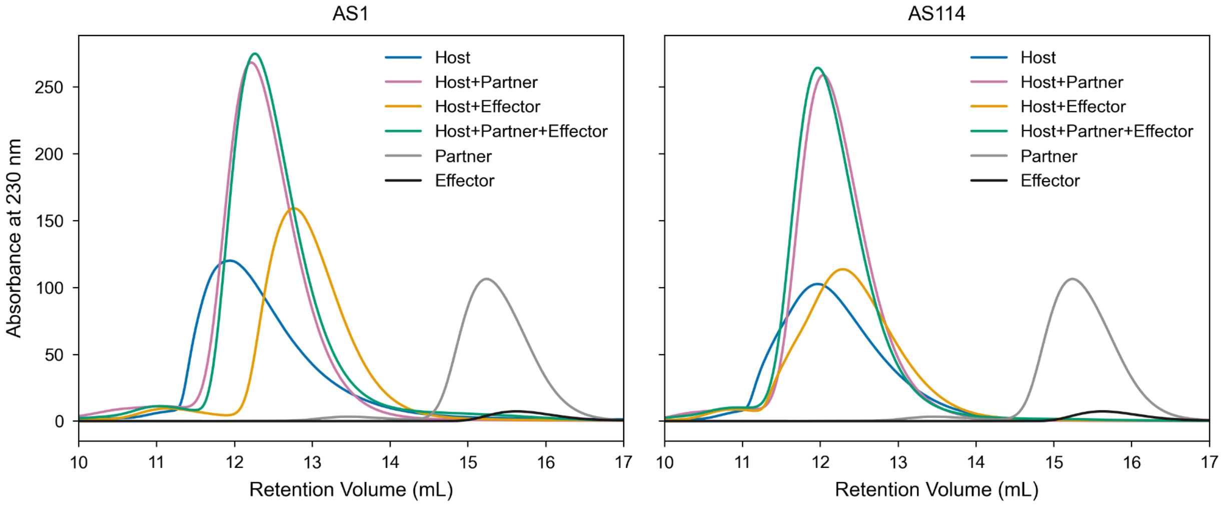


**Fig. S9: Stability of the AS1 and AS114 ternary complexes at high concentration.** SEC binding experiments performed on a Superdex 75 Increase 10/300 GL column. All components were injected at 20 µM. In isolation, the partner (gray) and effector (black) elute around 15–16 mL. When the host is included, depletion of signal in this range indicates partner-host or host-effector binding. The hosts alone (blue) run large and are likely weak homodimers. Including either partner (pink) or effector (orange) reduces homodimerization so the complex elutes later; signal past 15 mL is fully depleted, indicating complete binding to the host. Including both partner and effector (green) near-fully (for AS1) or fully (for AS114) depletes signal past 15 mL, indicating both partner and effector are bound to the host. SEC is a diluting and inherently nonequilibrium binding measurement: complexes which dissociate while on the column will elute later. This explains the long tail extending past 15 mL observed for the AS1 ternary complex. At the time of injection, the partner was likely completely bound, and since the partner dissociates from AS1 20-fold faster than from AS114 (Fig. 4a), some partner dissociated while on the column, leaving a tail. Thus, when both partner and effector are present at sufficiently high concentrations, the ternary complex is the dominant state.

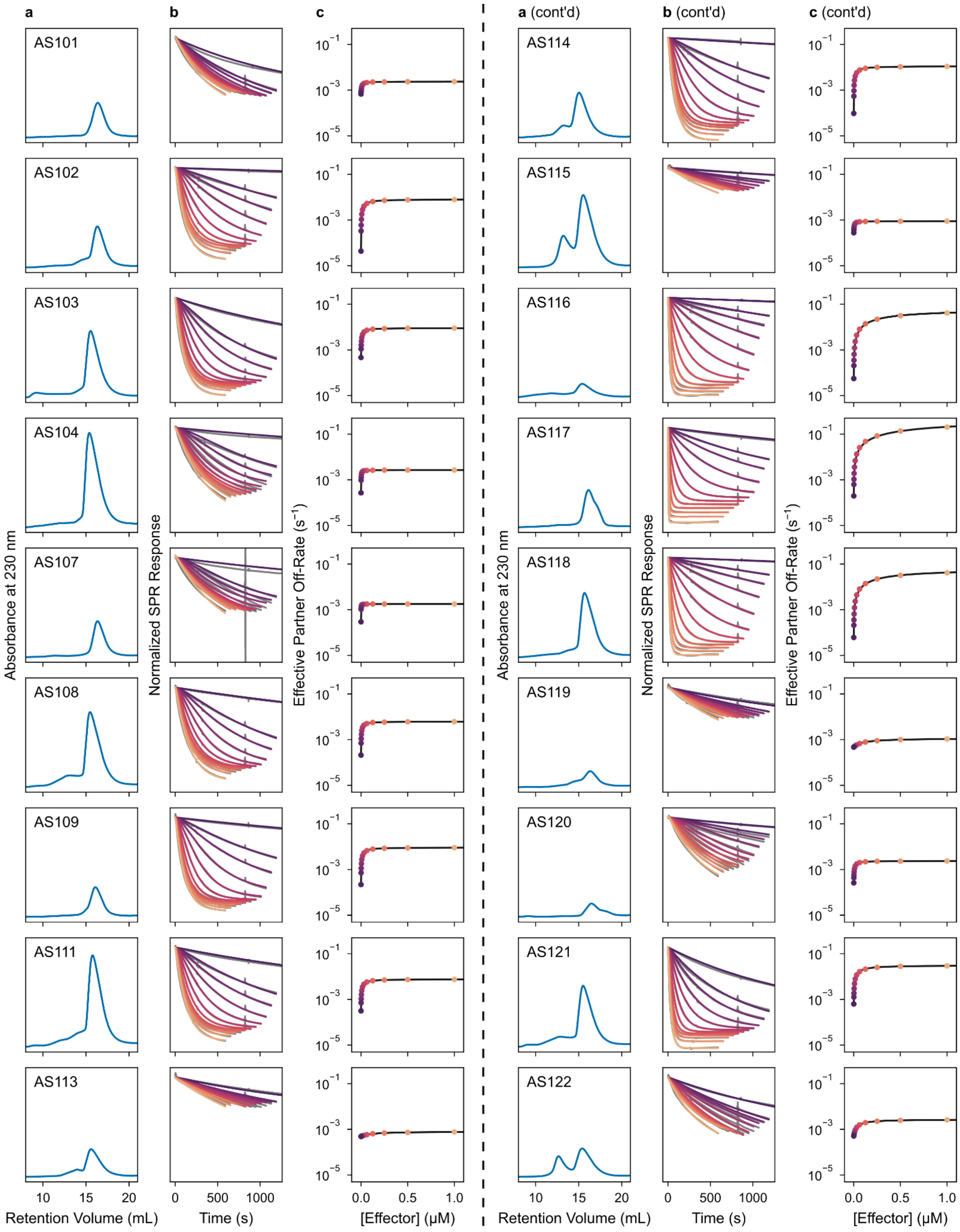
a
b
c
a (cont'd)
b (cont'd)
c (cont'd)
AS101
AS102
AS103
AS104
AS107
AS108
AS109
AS111
AS113
AS114
AS115
AS116
AS117
AS118
AS119
AS120
AS121
AS122
Absorbance at 230 nm
Normalized SPR Response
Effective Partner Off-Rate (s$^{-1}$)
$10^{-1}$
$10^{-3}$
$10^{-5}$
10
15
20
0
500
1000
0.0
0.5
1.0
Retention Volume (mL)
Time (s)
[Effector] (μM)

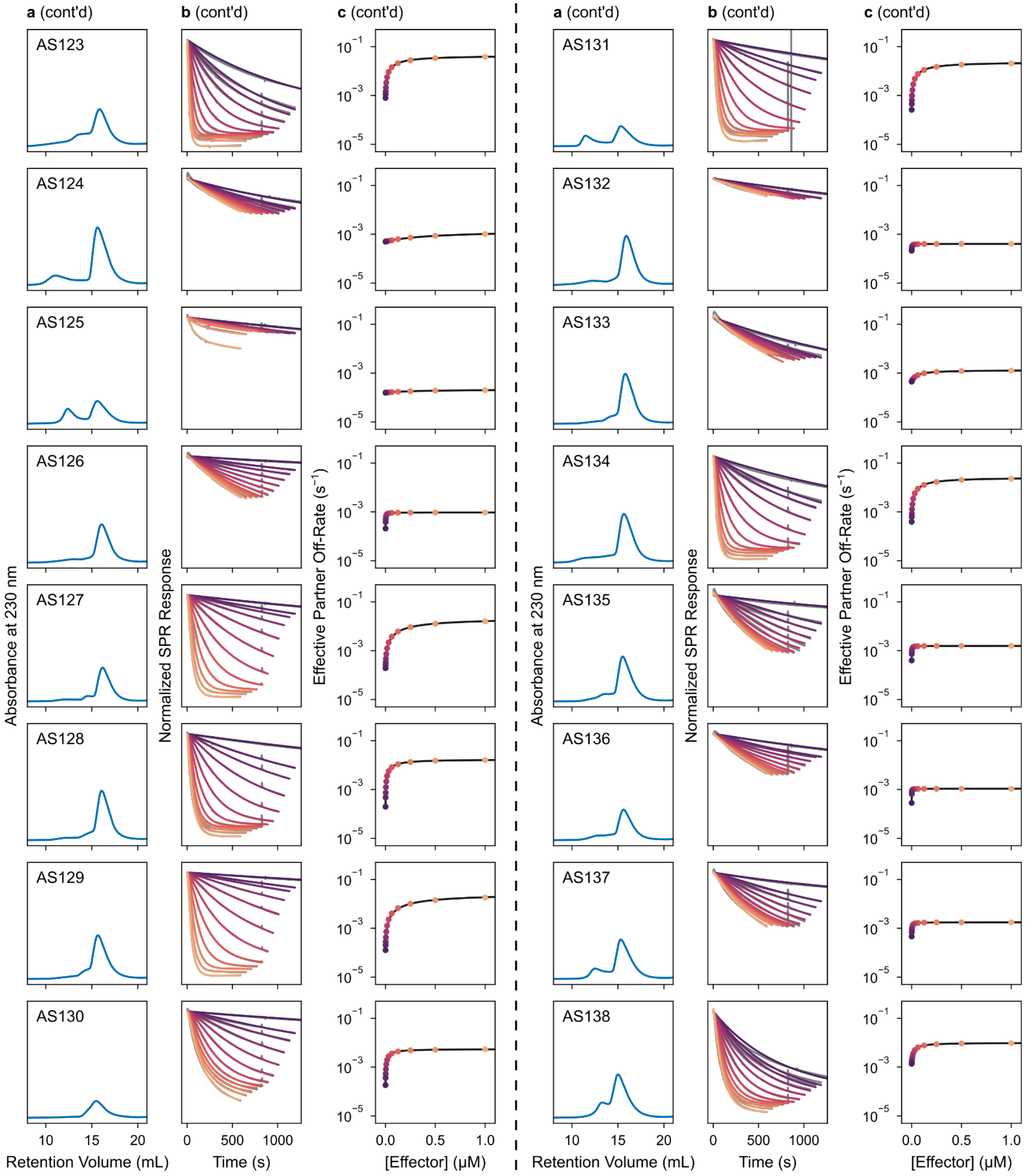


**Fig. S10: Characterization of AS1 variants. a**, SEC host purifications performed on a Superdex 200 Increase 10/300 GL column. **b**, Effector concentration–dependent dissociation of the partner upon addition of peptide effector. Data (gray) fit (colors) as described in methods. **c**, Effective partner off-rates computed from the model fit by ln(2)÷{half-time of partner-host interaction} plotted against effector concentration (circles) and fit with hyperbolic equations (black lines).

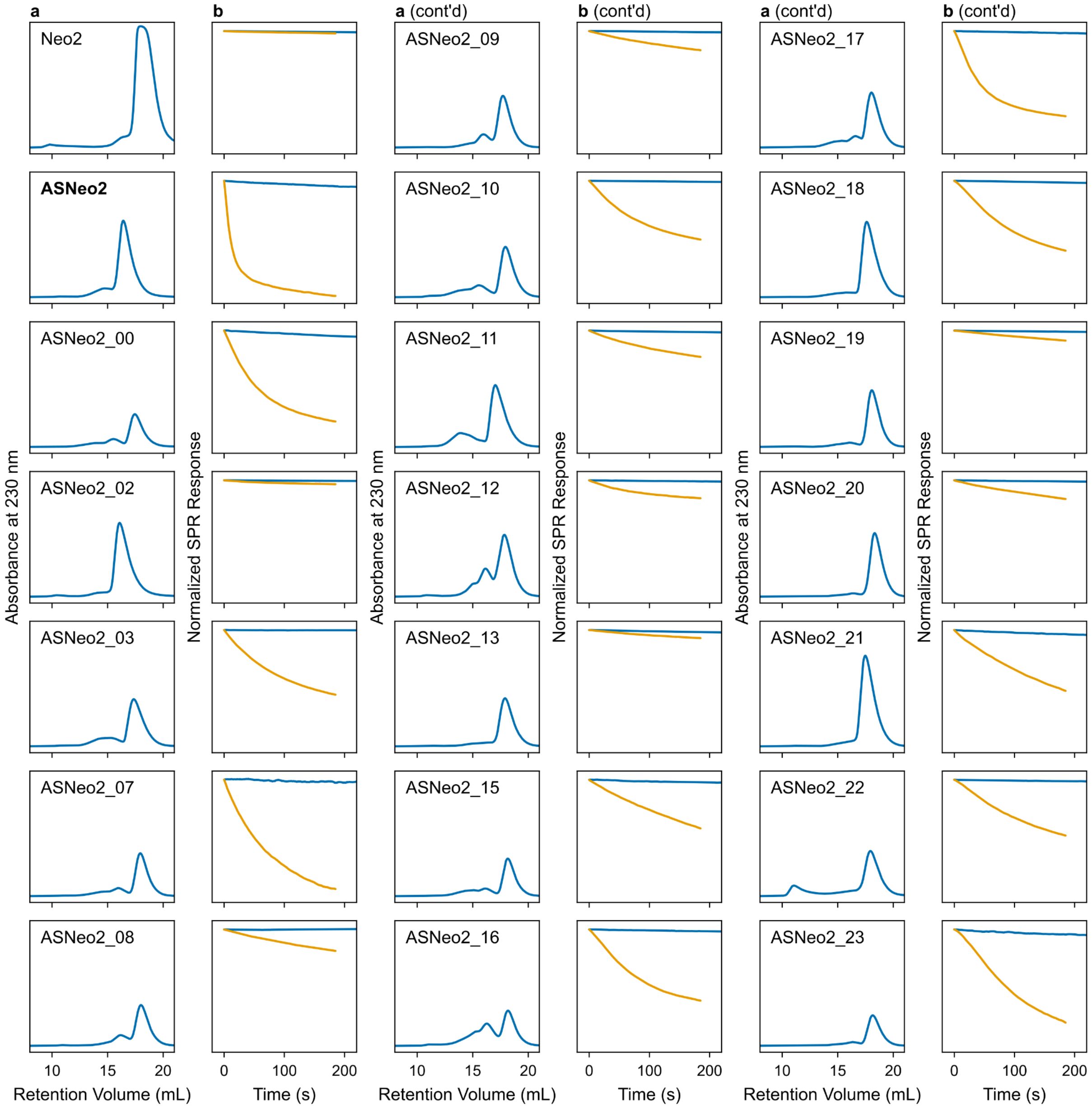


**Fig. S11: Characterization of initial switchable IL-2 mimic designs.** **a**, SEC purifications performed on a Superdex 200 Increase 10/300 GL column. **b**, Slow dissociation of $\gamma_c$ from the ASNeo2-IL-2R$\beta\gamma_c$ complex in the absence of effector (blue) and faster dissociation in the presence of effector (orange) as assessed by SPR.

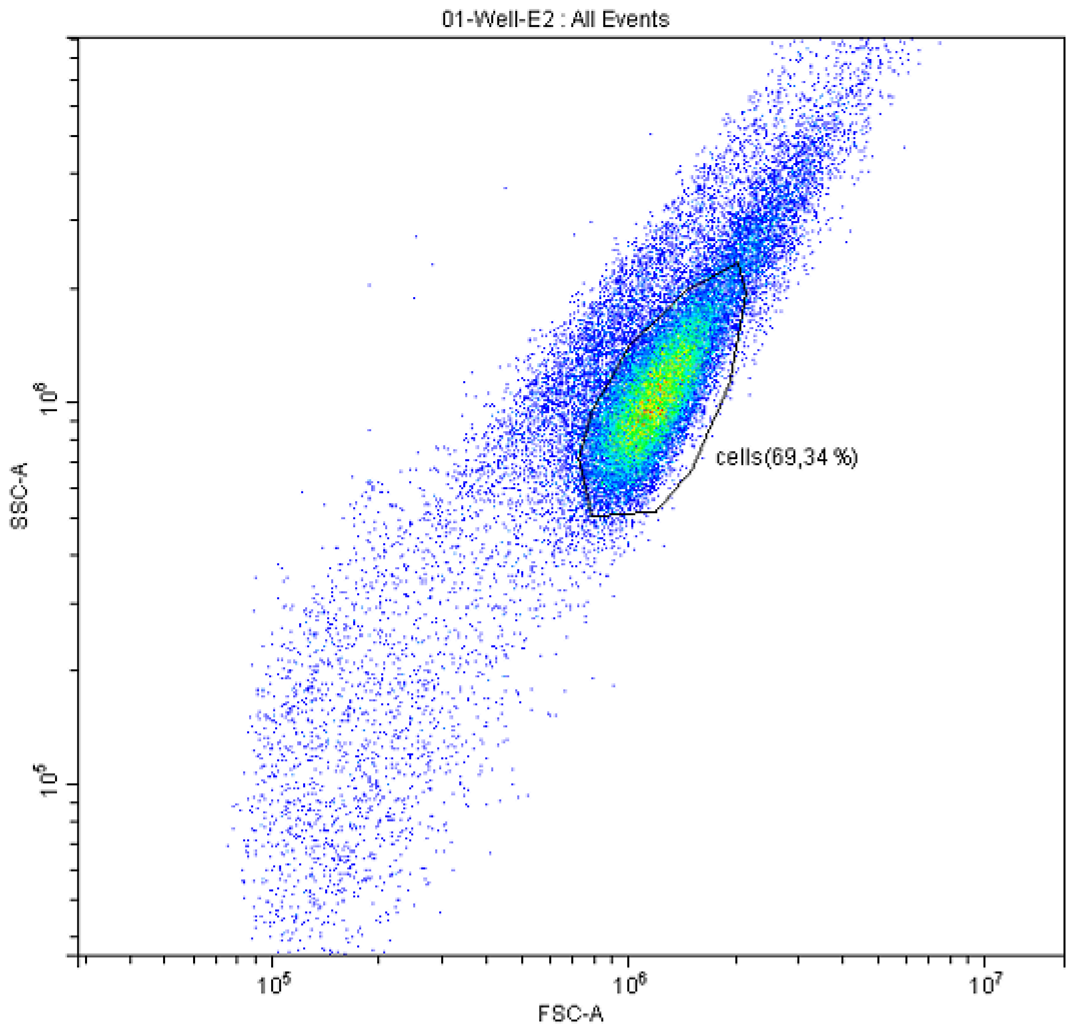


**Figure S12: Flow cytometry gating.** YT cell populations were identified based on forward/side scatter profiles.

**Supplementary Table 1: Kinetic parameters from facilitated dissociation experiments with peptide effector.**

| **Design** | $k_{off,B:A}$ | $k_{off,B:AC}$ | **Peptide Effector $EC_{50}$ (nM)** |
|---|---|---|---|
| LHD101B4 | 9e-5 | - | - |
| AS0 | 2e-4 | 1e-1 | 6,000 |
| AS1 | 2e-3 | 2e-1 | 100 |
| AS2 | 3e-4 | 4e-2 (2e-2) | 80 |
| AS5 | 4e-4 | 3e-2 | 10 |
| AS7 | 3e-3 | 8e-2 (3e-2) | 130 |
| AS101 | 7e-4 | 2e-3 | 5 |
| AS102 | 4e-5 | 8e-3 | 2 |
| AS103 | 5e-4 | 9e-3 | 4 |
| AS104 | 3e-4 | 3e-3 | 0.8 |
| AS107 | 3e-4 | 2e-3 | 0.4 |
| AS108 | 2e-4 | 6e-3 | 2 |
| AS109 | 2e-4 | 9e-3 | 3 |
| AS111 | 3e-4 | 8e-3 | 4 |
| AS113 | 5e-4 | 8e-4 | 100 |
| AS114 | 1e-4 | 1e-2 | 3 |
| AS115 | 3e-4 | 9e-4 | 3 |
| AS116 | 6e-5 | 6e-2 | 10 |
| AS117 | 2e-4 | 5e-1 | 30 |
| AS118 | 6e-5 | 6e-2 | 10 |
| AS119 | 5e-4 | 1e-3 | 90 |
| AS120 | 3e-4 | 2e-3 | 4 |
| AS121 | 6e-4 | 3e-2 | 8 |
| AS122 | 5e-4 | 3e-3 | 30 |
| AS123 | 8e-4 | 4e-2 | 20 |
| AS124 | 5e-4 | 1e-3 | 500 |

| | | | |
|---|---|---|---|
| AS125 | 2e-4 | 2e-4 | 300 |
| AS126 | 2e-4 | 9e-4 | 1 |
| AS127 | 2e-4 | 2e-2 | 30 |
| AS128 | 2e-4 | 2e-2 | 8 |
| AS129 | 1e-4 | 3e-2 | 30 |
| AS130 | 2e-4 | 5e-3 | 6 |
| AS131 | 3e-4 | 2e-2 | 20 |
| AS132 | 2e-4 | 4e-4 | 2 |
| AS133 | 4e-4 | 1e-3 | 50 |
| AS134 | 4e-4 | 3e-2 | 20 |
| AS135 | 4e-4 | 2e-3 | 1 |
| AS136 | 3e-4 | 1e-3 | 0.6 |
| AS137 | 5e-4 | 2e-3 | 2 |
| AS138 | 1e-3 | 1e-2 | 20 |
| ASNeo2 | 2e-4 | 2e-1 | 500 |
| ASNeo2_cp03 | 2e-4 | 9e-2 | 40 |
| ASNeo2_cp08 | 4e-5 | 2e-1 | 200 |
| ASNeo2_cp12 | 6e-5 | 2e-1 | 300 |

Values in parentheses give the maximum rate of partner dissociation induced by effector binding (the full facilitated dissociation pathway), if different from the value of $k_{off,B:AC}$ measured directly by forming the ternary complex on the SPR chip. The $EC_{50}$ is the concentration of effector required to achieve half the total acceleration of partner dissociation on a log scale.

**Supplementary Table 2: DEER experimental and fit parameters**

| Construct | Sites | State | $\tau_2$ (µs) | $\Delta t$ (ns) | Scans | $\lambda$ | SNR | $t_0$ offset (ns) | $\alpha$ |
|---|---|---|---|---|---|---|---|---|---|
| AS1 | R35R1<br>E173R1 | A | 6.000 | 22 | 47 | 0.49 | 117 | 84.6 | 23.89 |
| AS1 | R35R1<br>E173R1 | A+B | 6.000 | 22 | 53 | 0.45 | 47 | 113.9 | 36.92 |
| AS1 | R35R1<br>E173R1 | A+C | 6.000 | 22 | 85 | 0.46 | 87 | 98.4 | 40.00 |
| AS1 | R35R1<br>E173R1 | A+B+C | 6.000 | 22 | 135 | 0.42 | 46 | 124.2 | 10.71 |
| AS114 | E31R1<br>E165R1 | A | 7.000 | 22 | 27 | 0.47 | 32 | 91.5 | 50.00 |
| AS114 | E31R1<br>E165R1 | A+B | 6.000 | 22 | 93 | 0.38 | 60 | 79.4 | 6.46 |
| AS114 | E31R1<br>E165R1 | A+C | 6.000 | 22 | 140 | 0.42 | 89 | 91.5 | 7.06 |
| AS114 | E31R1<br>E165R1 | A+B+C | 6.000 | 22 | 138 | 0.43 | 70 | 86.3 | 22.06 |

$\Delta t$ - Pump pulse time step
$\lambda$ - Modulation depth
SNR - Signal-to-noise
$\alpha$ - Smoothing parameter

**Supplementary Table 3: Crystallographic data collection and refinement statistics.**

| | CS221B (PDB Code:9DD5) | LHD101An1 (PDB Code:9DD4) | AS5_AC (PDB Code:9DD3) | AS5_A (PDB Code:9DD2) |
|---|---|---|---|---|
| Resolution range | 33.81 - 1.5 (1.62 - 1.50) | 51.88 - 2.11 (2.17 - 2.11) | 31.69 - 1.64 (1.68 - 1.64) | 48.84 - 1.72 (1.75 - 1.72) |
| Space group | *C 2* | $P\ 2_1 2_1 2_1$ | *C 2* | *P 43* |
| Unit cell | 54.67, 25.09, 34.67; 90, 102.77, 90 | 64.97, 70.66, 76.39; 90, 90, 90 | 90.17, 35.66, 75.52; 90, 116.2, 90 | 62.37, 62.37, 78.50; 90, 90, 90 |
| Unique reflections | 7489 (1451) | 20777 (1711) | 24347 (1919) | 31981 (1667) |
| Multiplicity | 6.7 (6.8) | 6.4 (6.5) | 1.9 (1.9) | 9.0 (9.2) |
| Completeness (%) | 99.71 (99.38) | 99.63 (99.82) | 91.01 (95.00) | 100.00 (100.00) |
| Mean I/sigma(I) | 12.57 (1.71) | 21.25 (1.80) | 6.62 (1.14) | 18.50 (2.2) |
| Wilson B-factor | 22 | 62 | 28 | 32 |
| R-merge | 0.060 (0.788) | 0.037 (0.911) | 0.062 (0.537) | 0.047 (0.709) |
| R-pim | 0.025 (0.324) | 0.015 (0.384) | 0.054 (0.466) | 0.017 (0.263) |
| $CC_{1/2}$ | 0.999 (0.922) | 1.000 (0.929) | 0.993 (0.717) | 1.000 (0.875) |
| Reflections used in refinement | 7489 (1451) | 20777 (1711) | 24347 (1919) | 31945 (2281) |
| R-work | 0.2091 (0.2512) | 0.2222 (0.3295) | 0.2018 (0.3285) | 0.2035 (0.2755) |
| R-free | 0.2494 (0.3124) | 0.2617 (0.3301) | 0.2509 (0.3520) | 0.2431 (0.3195) |

| | | | | |
|---|---|---|---|---|
| Number of non-hydrogen atoms | 475 | 1992 | 2219 | 2126 |
| macromolecules | 430 | 1976 | 2149 | 2001 |
| solvent | 45 | 16 | 70 | 125 |
| Protein residues | 52 | 246 | 278 | 258 |
| RMS(bonds) | 0.004 | 0.002 | 0.003 | 0.006 |
| RMS(angles) | 0.60 | 0.35 | 0.53 | 0.68 |
| Ramachandran favored (%) | 100.00 | 97.93 | 98.18 | 99.22 |
| Ramachandran allowed (%) | 0.00 | 2.07 | 1.82 | 0.78 |
| Ramachandran outliers (%) | 0.00 | 0.00 | 0.00 | 0.00 |
| Average B-factor | 32 | 80 | 35 | 41 |
| macromolecules | 31 | 80 | 35 | 40 |
| solvent | 46 | 69 | 41 | 46 |

The highest-resolution shell are shown in parentheses.

| | AS1_A (PDB Code:9DCX) | AS1_AC (PDB Code:9DCY) | AS1_ABC (PDB Code:9DD1) | AS1_AB #2 (PDB Code:9DD0) | AS1_AB #1 (PDB Code:9DCZ) |
|---|---|---|---|---|---|
| Resolution range | 48.58 - 1.80 (1.85 - 1.80) | 47.98 - 2.35 (2.47 - 2.35) | 28.21 - 3.70 (3.93 - 3.70) | 30.25 - 3.88 (4.34 - 3.88) | 34.75 - 2.90 (2.94 - 2.90) |
| Space group | *P 61 2 2* | $P\ 2_1$ | $P\ 43\ 2_1\ 2$ | $P\ 2_1 2_1 2_1$ | $P\ 6_1\ 2\ 2$ |
| Unit cell | 61.54, 61.54, 236.10; 90, 90, 120 | 78.44, 35.98, 88.90; 90. 109.63, 90 | 110.85, 110.85, 65.53; 90, 90, 90 | 62.98, 74.07, 211.77; 90, 90, 90 | 80.71, 80.71, 546.85; 90, 90, 120 |
| Unique reflections | 25499 (1764) | 19956 (2831) | 10334 (1717) | 9716 (2690) | 25024 (3898) |
| Multiplicity | 12.7 (13.3) | 4.8 (5.1) | 11.8 (12.1) | 9.6 (9.4) | 31.4 (34.2) |
| Completeness (%) | 99.31 (100.00) | 99.66 (99.89) | 100.00 (100.00) | 99.80 (100.00) | 99.90 (100) |
| Mean I/sigma(I) | 22.53 (8.59) | 8.70 (1.88) | 10.70 (2.65) | 5.80 (3.5) | 10.5 (2.1) |
| Wilson B-factor | 33 | 31 | 103 | 122 | 66 |
| R-merge | 0.084 (0.945) | 0.090 (0.221) | 0.190 (0.951) | 0.270 (0.717) | 0.369 (2.128) |
| R-pim | 0.018 (0.246) | 0.044 (0.109) | 0.057 (0.280) | 0.091 (0.246) | 0.067 (0.369) |
| $CC_{1/2}$ | 0.955 (0.673) | 0.997 (0.958) | 0.997 (0.692) | 0.994 (0.892) | 0.998 (0.876) |
| Reflections | 25499 (1764) | 19956 (2831) | 4667 (756) | 9667 (1344) | 24845 (960) |

| | | | | | |
|---|---|---|---|---|---|
| used in refinement | | | | | |
| R-work | 0.2135 (0.2862) | 0.1937 (0.2226) | 0.2472 (0.3015) | 0.2261 (0.2743) | 0.2094 (0.3779) |
| R-free | 0.2552 (0.3298) | 0.2437 (0.3186) | 0.2772 (0.3010) | 0.2789 (0.3296) | 0.2550 (0.4580) |
| Number of non-hydrogen atoms | 2059 | 4409 | 3062 | 5891 | 6025 |
| macromolecules | 1965 | 4340 | 3062 | 5891 | 6010 |
| solvent | 92 | 59 | n/a | n/a | 7 |
| Protein residues | 257 | 564 | 390 | 749 | 759 |
| RMS(bonds) | 0.008 | 0.001 | 0.002 | 0.003 | 0.003 |
| RMS(angles) | 0.81 | 0.34 | 0.40 | 0.53 | 0.520 |
| Ramachandran favored (%) | 99.22 | 98.02 | 96.59 | 97.46 | 96.35 |
| Ramachandran allowed (%) | 0.78 | 1.98 | 3.41 | 2.37 | 3.48 |
| Ramachandran | 0.00 | 0.00 | 0.00 | 0.17 | 0.17 |

| | | | | | |
|---|---|---|---|---|---|
| outliers (%) | | | | | |
| Average B-factor | 39 | 42 | 92 | 115 | 84 |
| macromolecules | 38 | 42 | 92 | 115 | 84 |
| solvent | 44 | 37 | n/a | n/a | 56 |

The highest-resolution shell are shown in parentheses.
